# Cerebellar Purkinje cells can differentially modulate coherence between sensory and motor cortex depending on region and behavior

**DOI:** 10.1101/2020.03.11.986943

**Authors:** Sander Lindeman, Lieke Kros, Sungho Hong, Jorge F. Mejias, Vincenzo Romano, Mario Negrello, Laurens W.J. Bosman, Chris I. De Zeeuw

## Abstract

Coherence among sensory and motor cortices is indicative of binding of critical functions in perception, motor planning, action and sleep. Evidence is emerging that the cerebellum can impose coherence between cortical areas, but how and when it does so is unclear. Here, we studied coherence between primary somatosensory (S1) and motor (M1) cortices during sensory stimulation of the whiskers in the presence and absence of optogenetic stimulation of cerebellar Purkinje cells in awake mice. Purkinje cell activation enhanced and reduced sensory-induced S1-M1 coherence in the theta and gamma bands, respectively. This impact only occurred when Purkinje cell stimulation was given simultaneously with sensory stimulation; a 20 ms delay was sufficient to alleviate its impact, suggesting the existence of a fast, cerebellar sensory pathway to S1 and M1. The suppression of gamma band coherence upon Purkinje cell stimulation was significantly stronger during trials with relatively large whisker movements, whereas the theta band changes did not show this correlation. In line with the anatomical distribution of the simple spike and complex spike responses to whisker stimulation, this suppression also occurred following focal stimulation of medial crus 2, but not of lateral crus 1. Granger causality analyses and computational modeling of the involved networks suggest that Purkinje cells control S1-M1 coherence most prominently via the ventrolateral thalamus and M1. Our results indicate that coherences between sensory and motor cortices in different frequency ranges can be dynamically modulated by cerebellar input, and that the modulation depends on the behavioral context and is site-specific.

**Significance Statement:** Coherent activity between sensory and motor areas is essential in sensorimotor integration. We show here that the cerebellum can differentially affect cortical theta and gamma band coherences evoked by whisker stimulation via a fast ascending and predictive pathway. In line with the functional heterogeneity of its modular organization, the impact of the cerebellum is region-specific and tuned to ongoing motor responses. These data highlight site-specific and context-dependent interactions between the cerebellum and the cerebral cortex that can come into play during a plethora of sensorimotor functions.

## Introduction

Coherent oscillations can bind different brain areas by affecting susceptibility of neurons to synaptic input and providing a timing mechanism for generating a common dynamical frame for cortical operations (Engel et al., 1992; Fries, 2015). For example, coherence can create a temporal framework for concerted neural activity that facilitates integration of the activity of sensory and motor areas (Bressler et al., 1993; Ahrens and Kleinfeld, 2004; Womelsdorf and Fries, 2006). Online integration is particularly relevant, when animals, including ourselves, explore their environment via active touch, requiring sensory input to be directly related to the momentary position and movement of eyes, fingertips, antennae, whiskers or other organs (Bosman et al., 2011; Hartmann, 2011; Prescott et al., 2011).

Coherence often occurs in specific frequency bands that can be associated with different functions. In the field of sensorimotor integration, skilled movements rely on intercortical coherence between sensory and motor areas that occur in the theta (4-8 Hz) range during force generation, while coherence at higher bands is engaged during the preparation thereof (Arce-McShane et al., 2016). Likewise, within the field of visual perception, coherence in the alpha (8-12 Hz) and gamma (30-100 Hz) bands have been found to contribute to feedback and feedforward processing, respectively (van Kerkoerle et al., 2014; Shin and Moore, 2019).

The appearance of coherence among different cortical regions implicates reciprocal connections between neurons distributed among different layers within the cerebral cortex (Cardin et al., 2009; Buzsáki and Wang, 2012; Jensen et al., 2014; Mejias et al., 2016; Veit et al., 2017) as well as inputs from subcortical structures like the thalamus (Pedroarena and Llinás, 1997; Saalmann et al., 2012; Song and Francis, 2015; Jaramillo et al., 2019) (Fig. 1A). Accordingly, one of the main inputs to the thalamus, i.e., the cerebellum, has a strong impact in organizing cortico-cortical coherence (O’Connor et al., 2002; Popa et al., 2013; Vecchio et al., 2019). Indeed, disruption of cerebellar function, whether inflicted pharmacologically in rats (Popa et al., 2013) or due to stroke in patients (Vecchio et al., 2019), affects cortico-cortical coherence.

**Figure 1.**
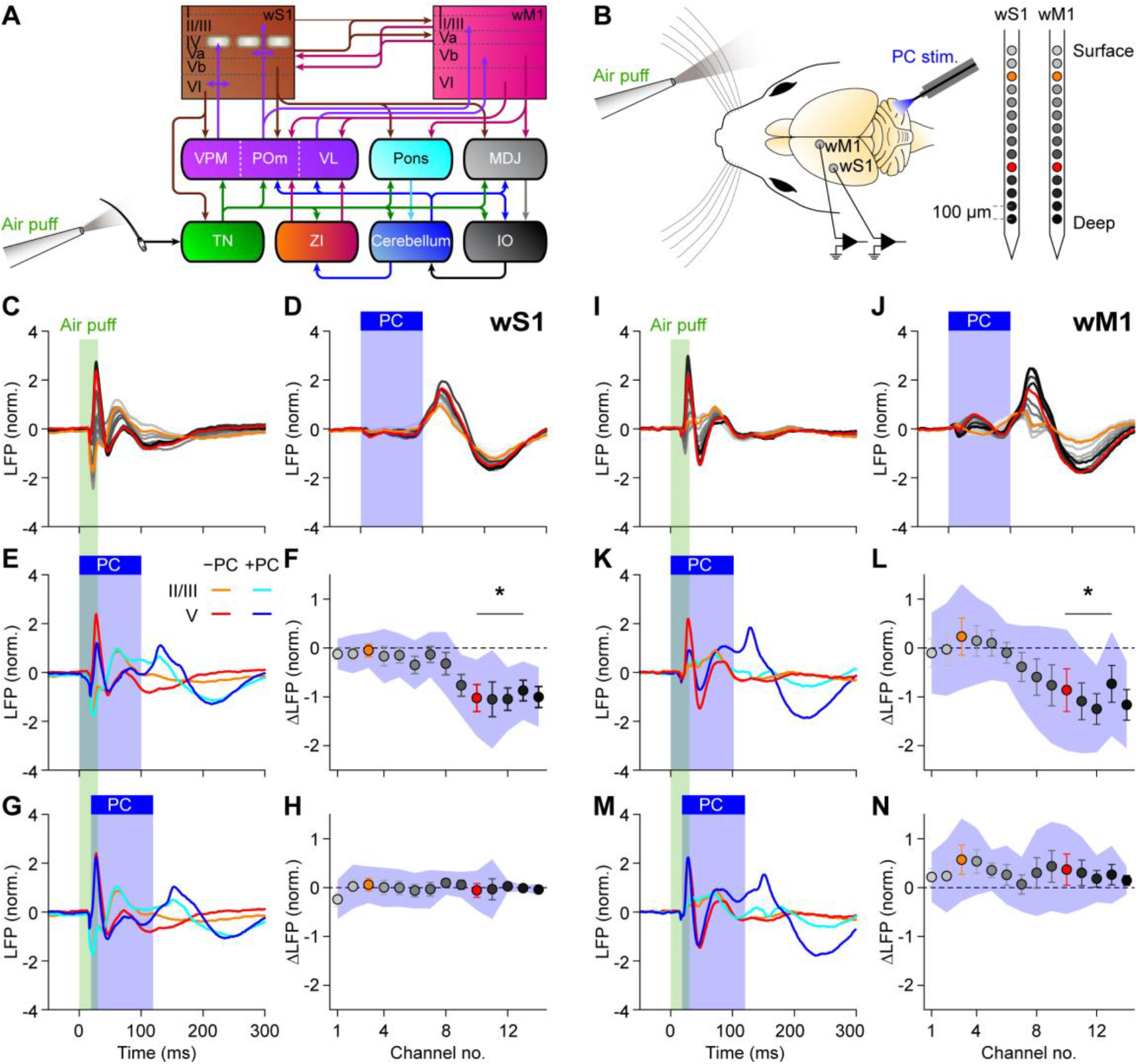
Purkinje cell stimulation disinhibits fast sensory responses in wS1 and wM1. **A** Simplified scheme of anatomical pathways carrying whisker input to wS1, wM1 and the cerebellum, and the reciprocal cerebro-cerebral connections. IO = inferior olive; MDJ = nuclei of the mesodiencephalic junction; Pom = posterior medial nucleus; TN = trigeminal nuclei; VL = ventrolateral nucleus; VPM = ventroposterior medial nucleus; ZI = zona incerta. **B** Local field potentials (LFP) were recorded in wS1 and wM1 of awake mice using, for each area, 14 recording spots on linear silicon probes. Colors indicate their relative positions, with orange and red for the 3^rd^ and 10^th^ electrodes, representing the supra- and subgranular layers, respectively. Purkinje cells (PC) were stimulated optogenetically using an optic fiber with 400 μm diameter placed on the center of crus 1 (Fig. S1). **C** Whisker stimulation triggered fast responses in contralateral wS1, as illustrated by the averaged LFP traces (*n* = 100 trials per mouse, *N* = 8 mice). **D** Purkinje cell stimulation triggered delayed responses in wS1 after rebound firing in the cerebellar nuclei (Fig. S2B). **E** Comparison of the LFPs recorded during trials with whisker stimulation (orange / red) and with combined with sensory and Purkinje cell stimulation (cyan / blue). **F** During the early response period, especially the amplitude of the first positive LFP peak in the subgranular layers was affected, in addition to profound impact during later phases of sensory processing. Plotted are the averaged differences in amplitude of the first positive LFP peaks. Error bars indicate SEM and shaded area sd. **G**-**H** The impact of optogenetic Purkinje cell stimulation on the first positive LFP peak was largely abolished by introducing a 20 ms delay between the start of air puff sensory stimulation and the onset optogenetic Purkinje cell stimulation. **I**-**N** The same plots as **C**-**H**, but now for wM1, showing comparable impact of optogenetic Purkinje stimulation on the sensory-induced LFP signals.

Even though the impact of cerebellar activity on cortical coherence is well established (Popa et al., 2013; Proville et al., 2014; Kros et al., 2015), it remains to be elucidated to what extent and how the cerebellum can differentially influence different frequency bands, to what extent such potentially different impacts depend on the behavioral context, and whether these differential effects are mediated through different cerebellar modules (Proske and Gandevia, 2009; De Zeeuw et al., 2011; Romano et al., 2018). Here, we set out to address these questions by investigating the impact of Purkinje cell activity on coherence between the whisker areas of the primary somatosensory (wS1) and motor cortex (wM1) during stimulation of the whiskers in awake behaving mice. When we stimulated Purkinje cells optogenetically at different intervals with respect to air puff stimulation of the whiskers, we observed that these main output neurons differentially contribute to wS1-wM1 theta and gamma band coherences with opposite effects, depending on ongoing behavior and their precise site in the cerebellar hemispheres.

## Results

### Purkinje cell stimulation modulates sensory responses in wS1 and wM1

Whisker stimulation triggers fast responses in wS1 and wM1 (Kleinfeld et al., 2002; De Kock et al., 2007; Ferezou et al., 2007) as well as in the cerebellar cortex (Bosman et al., 2010; Brown and Raman, 2018; Romano et al., 2018). Before studying the impact of cerebellar stimulation on the cortical coherence between wS1 and wM1, we first needed to determine to what extent the individual cortical responses within wS1 and wM1 depended on cerebellar activity.

Thereto, we compared local field potentials (LFPs) in wS1 during whisker stimulation in the absence or presence of optogenetic stimulation of Purkinje cells in the crus 1 and crus 2 area (Figs. 1B, S1). Air puffs applied to the whiskers evoked canonical LFP responses in wS1 in that they induced an initial fast decrease, first in layer IV and then also in the superficial and deep layers, followed by increased LFP signals (Figs. 1C, S2A, S3A). Decreases is LFP signal have been suggested to correspond with increased neuronal excitation (Pettersen et al., 2008), returned to baseline after roughly 200 ms.

Given the strong and direct trigemino-thalamo-cortical pathways (Yu et al., 2006; Bosman et al., 2011), a significant impact of Purkinje cell activation during whisker stimulation on the initial response in the input layer of wS1 would not be expected. Indeed, this was not the case (*p* = 0.197, Fig. S4A; see Table S1 for details on statistical analysis). However, the spread of excitation to the deeper layers was enhanced by Purkinje cell stimulation (*p* = 0.012, Fig. S4A). Likewise, Purkinje cell stimulation reduced the positive LFP peak in the deeper layers (*p* = 0.024, Figs. 1E-F, S2A, S3A). Both effects were not observed when we postponed the Purkinje cell stimulation 20 ms relative to the whisker stimulation (Figs. 1G-H, S2A, S3A, S4A, Table S1). When we, as a control, stimulated the Purkinje cells in the absence of sensory stimulation, we observed that this induced a near-complete block of the output of cerebellar nucleus neurons, followed by rebound firing at the end of the 100 ms stimulus interval (Fig. S2B). Of note, the period of silencing led to a small, but observable increase in neural activity in wS1 (negative LFP) while the rebound firing in the cerebellar nuclei correlated with decreased neural activity in wS1 (positive LFP; Fig. 1D), suggesting that the connection between cerebellar nuclei and wS1 includes at least one inhibitory hub.

Similar to wS1, wM1 did not display a significant impact of Purkinje cell stimulation on the initial wave of neural excitation following sensory stimulation (Fig. S4B, Table S1). However, the subsequent positive peak in the LFP was reduced in the deeper layers (*p* = 0.024, Figs. 1I-L, S2A, S3B). As for wS1, this effect was abolished by a 20 ms delay between whisker and Purkinje cell simulation (Fig. 1M-N). Thus, our data indicate a fast ascending pathway via the cerebellum disinhibiting sensory responses in the deeper layers of wS1 and slightly later also in wM1.

### Cerebellar output differentially modulates S1-M1 coherence in theta and gamma bands

As we found cerebellar activity to be able to modulate early-phase sensory responses in both wS1 and wM1, we surmised that cerebellar activity could also affect sensory-related coherent activity between these areas. As there are particularly strong connections between the subgranular layers of wS1 and the supragranular layers of wM1 (Ahrens and Kleinfeld, 2004; Mao et al., 2011) (Fig. 1A), we initially focused on the coherence between these layers. Air puff stimulation of the whiskers triggered a fast increase in S1-M1 coherence, particularly in the beta and lower gamma band range, and to a lesser extent in the theta range (Fig. 2). Instead, whereas sensory stimulation combined with simultaneous optogenetic Purkinje cell stimulation led to a further enhancement in the sensory-induced coherence at the theta band, the same combination prominently reduced the sensory-induced coherence at the gamma band (Fig. 2A,B,E). Both of these modifications could be alleviated by delaying the optogenetic stimulation with 20 ms relative to the onset of whisker stimulation (Fig. 2A,B,E). The fact that a 20 ms lag between sensory and Purkinje cell stimulation was sufficient to, at least in part, rescue the original amplitude of sensory-evoked signals corroborates the notion that under normal physiological circumstances cerebellar modulation of cerebral coherence is probably mediated by a fast pathway. Granger causality can measure how much of the wM1 signal in a certain frequency band is controlled by the wS1 and vice versa. Applying that analysis on the sensory-induced coherence revealed that, for the comparison of the deep layers of S1 and the superficial layers of M1, both areas were involved in generation of the coherence (Fig. 2C-D).

**Figure 2.**
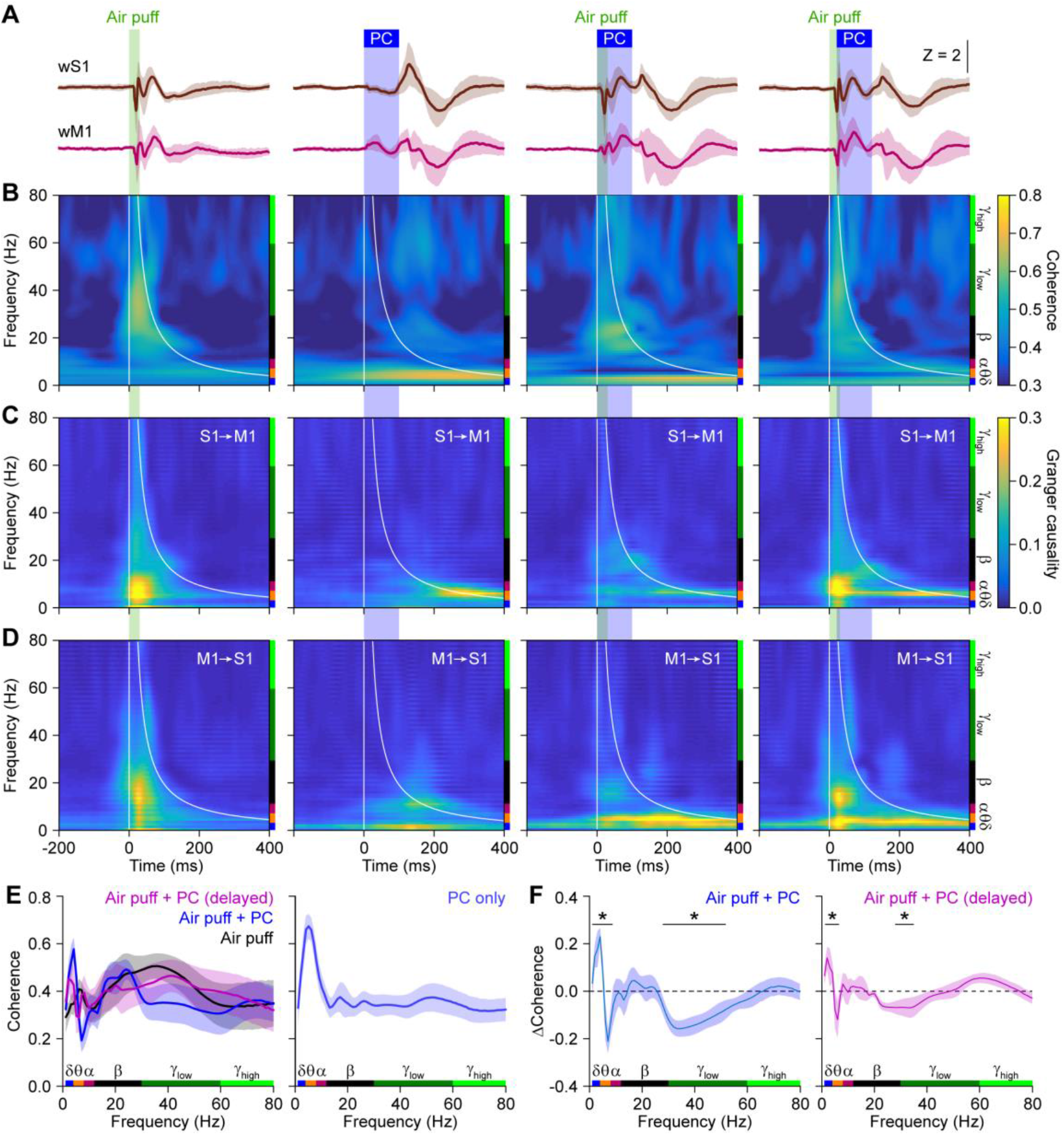
Reducing cerebellar output enhances and inhibits sensory-induced S1-M1 theta and gamma band coherence, respectively. **A** Averaged LFP signals in subgranular wS1 and supragranular wM1 following either air puff stimulation of the contralateral facial whiskers, optogenetic Purkinje cell (PC) stimulation, or a combination of both. In the column on the right, there was a 20 ms delay between the onset of air puff and Purkinje cell stimulation. Purkinje cell stimulation was performed with an optic fiber with a 400 μm diameter placed on the center of crus 1. **B** Heat maps with the coherence strength for each frequency. Purkinje cell stimulation induced a delayed increase in the lower frequency range, mainly theta band. Sensory stimulation predominantly caused a rapid increase in the lower gamma band range. This increased coherence in the lower gamma band range could be suppressed by simultaneous optogenetic stimulation of Purkinje cells. This suppression was largely absent when the optogenetic stimuli were delayed by 20 ms, indicating the importance of fast cerebellar processing. **C** Heat maps showing Granger causality from wS1 to wM1. **D** Granger causality for wM1 to wS1. **E** Coherence after whisker stimulation (left) and for optogenetic Purkinje cell stimulation alone (right). **F** Simultaneous Purkinje cell stimulation enhanced and suppressed theta and gamma band coherence, respectively. These modulations were largely reduced by a 20 ms delay in the onset of Purkinje cell stimulation. The increased theta band activity may partly reflect a direct effect of Purkinje cell stimulation. Shaded areas indicate SEM. *N =* 8 mice; * *p* < 0.05 (*χ^2^* > 3.84; difference of coherence test, see Methods)

Examining the coherence between other cortical layers, which are probably less directly coupled (Fig. 1A), we observed that the cerebellar impact broadened, now also comprising beta and higher gamma bands (Fig. S5). Granger causality analysis revealed that the sensory-induced gamma band coherence is largely triggered by the superficial layers of wS1 and from there spreads to the deep layers of wS1 (Fig. S6A-B) and the deep layers of wM1 (Fig. S6C-D). Notably, there was more balance between the deep layers of wS1 and the superficial layers of wM1 (Figs. 2C-D and S6C-D), suggesting that sensory-induced coherence is a complex phenomenon involving reciprocal connections between wS1 and wM1. The Granger causality analysis of the Purkinje cell-induced theta band coherence did not systematically reveal a strict directionality, suggesting a reciprocal involvement of wS1 and wM1.

Thus, experimentally dampening the output of the cerebellar nuclei by enhancing Purkinje cell activity results in an array of changes in sensory-induced coherences between wS1 and wM1. Effects were detected in all layers, but the most specific and reproducible changes were revealed in the comparison between theta and gamma coherence across wS1 and wM1.

### Cerebellar impact on S1-M1 coherence depends on behavioral context

Air puff stimulation of the whiskers triggers reflexive protraction, the amplitude of which is correlated to cerebellar activity (Brown and Raman, 2018; Romano et al., 2018). This suggests an interaction between cerebellar activity, whisker protraction and wS1-wM1 coherence. To sharpen this statement, we singled out, for each experiment, the 50% of the trials with the largest reflexive whisker protractions and compared these to the other 50% (Fig. 3A-B). During air puff stimulation, larger protractions were correlated to lower coherence levels than smaller protractions, which was opposite during combined whisker and Purkinje cell stimulation (Fig. 3C), alluding to a significantly stronger impact o f cerebellar activity during larger movements (Fig. 3D-E). Again, this impact was absent when introducing a 20 ms delay between whisker and Purkinje cell stimulation (Fig. 3D). The cerebellum therefore seems to provide contextual input with a homeostatic effect, which was particularly clear in the beta and gamma bands, but absent in the theta band. Purkinje cell stimulation in the absence of whisker stimulation did not cause a whisker movement until the end of the stimulus, as described previously (Proville et al., 2014) (Fig. S7).

**Figure 3.**
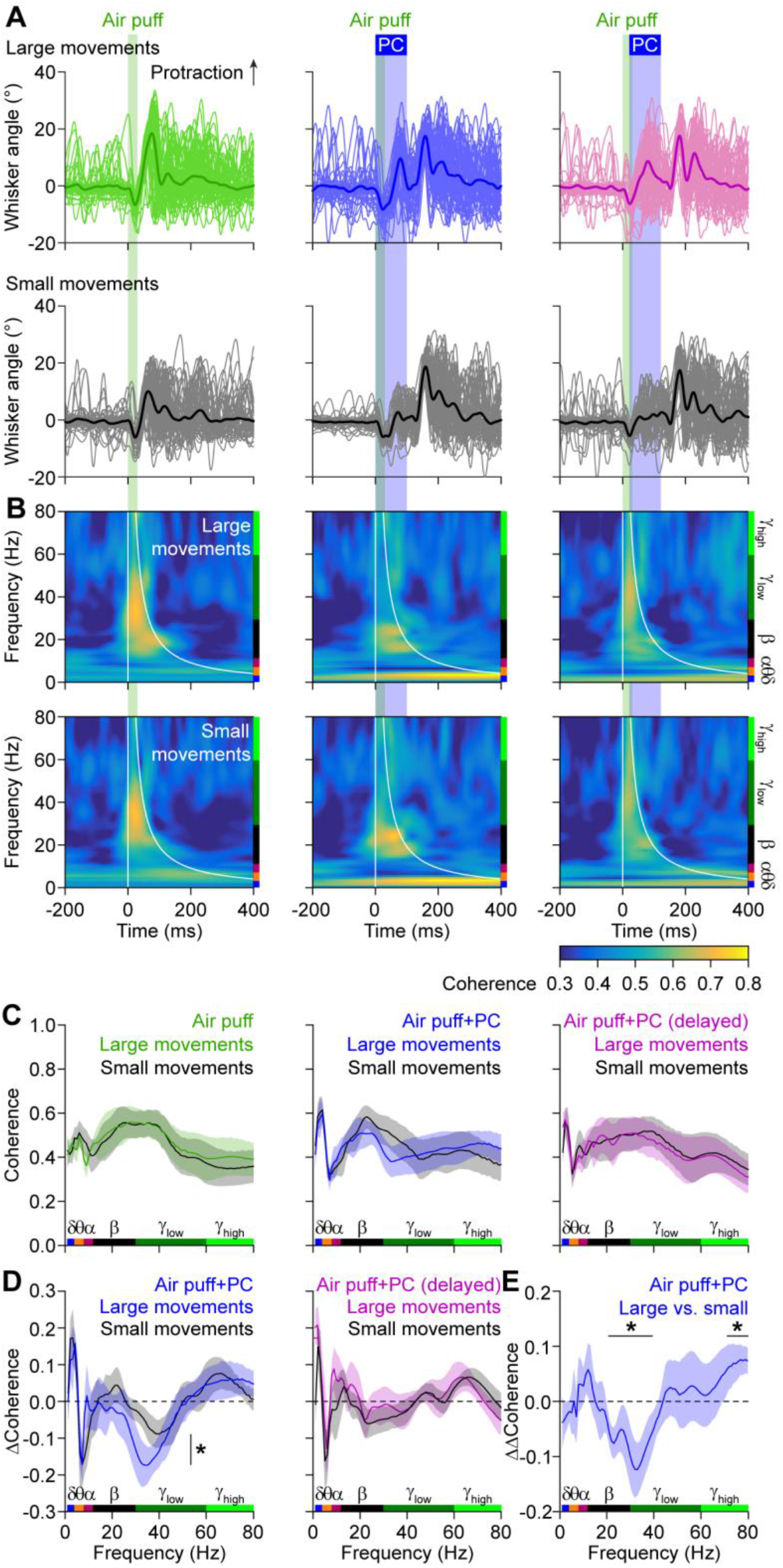
Cerebellar impact on wS1-wM1 coherence depends on behavior. **A** Whisker stimulation triggers reflexive protraction. Trials were split between the 50% of the trials with the largest and the 50% with the smallest protractions. **B** The coherence between the subgranular layers of wS1 and the subgranular layers of wM1 were only mildly different between the trials with large (upper row) and small (bottom row) movements. **C** For each stimulus condition, the averaged coherence spectra are plotted, with colored traces representing the large whisker movements. Note that the difference in beta and lower gamma band activity are modulated in opposite fashion when adding optogenetic Purkinje cell stimulation to the air puff stimulation. **D** Accordingly, the impact of optogenetic Purkinje cell stimulation on sensory-induced wS1-wM1 coherence was stronger during trials with large whisker movements. This difference was statistically significant (DoC test, see **E**). This effect was abolished by a 20 ms delay between the start of whisker and Purkinje cell stimulation. **E** The difference in the impact of simultaneous Purkinje cell stimulation (“DDCoherence”) on sensory-induced beta and gamma band coherence was significantly larger during the trials with large movements than during those with small movements (DoC analysis). Line in C - E indicate averages and the shades SEM. See also Fig. S7.

### Regional heterogeneity in cerebello-cerebral communication

The direction of Purkinje cell modulation upon whisker stimulation is related to the location within crus 1 and crus 2 receiving the stimulus (Romano et al., 2018). More specifically, whereas the increase in simple spike firing is most prominent in medial crus 2, that of the complex spikes, which may facilitate execution of touch-induced whisker protraction, is more robust in lateral crus 1 (Romano et al., 2018). Given this differential distribution in whisker-related Purkinje cell activity (Fig. 4A), we hypothesized that the changes in coherence described above depend on the specific area of optogenetic stimulation. To study the impact of spatial Purkinje cell stimulation on wS1 and wM1, we placed small optic fibers in a rectangular grid, targeting the medial and more lateral parts of crus 1 and crus 2. With these fibers we could reliably trigger a near-complete block of neuronal activity in the cerebellar nuclei and trigger whisker movements related to the rebound firing in these nuclei (Figs. S8 and S9). Yet, the illuminated volumes were small enough to reduce crosstalk between medial and lateral stimulus locations (Fig. S10). Consistent with the hypothesis described above, we found the most prominent differences in the impact of optogenetic stimulation during whisker stimulation between medial crus 2 and lateral crus 1 (Figs. 4B and S11). More specifically, comparison of the impact of Purkinje cell stimulation on the extent of sensory-induced coherence revealed that Purkinje cells in medial crus 2 and lateral crus 1 differed in their impact on gamma band coherence, but showed no significant difference on theta band coherence. Since these data on regional heterogeneity were based on differential Purkinje cell modulations during different forms of adaptive and reflexive whisking behavior (Romano et al., 2018), they are consistent with the prominent dependency of the gamma, but not theta, band coherence on behavioral context (Fig. 3).

**Figure 4.**
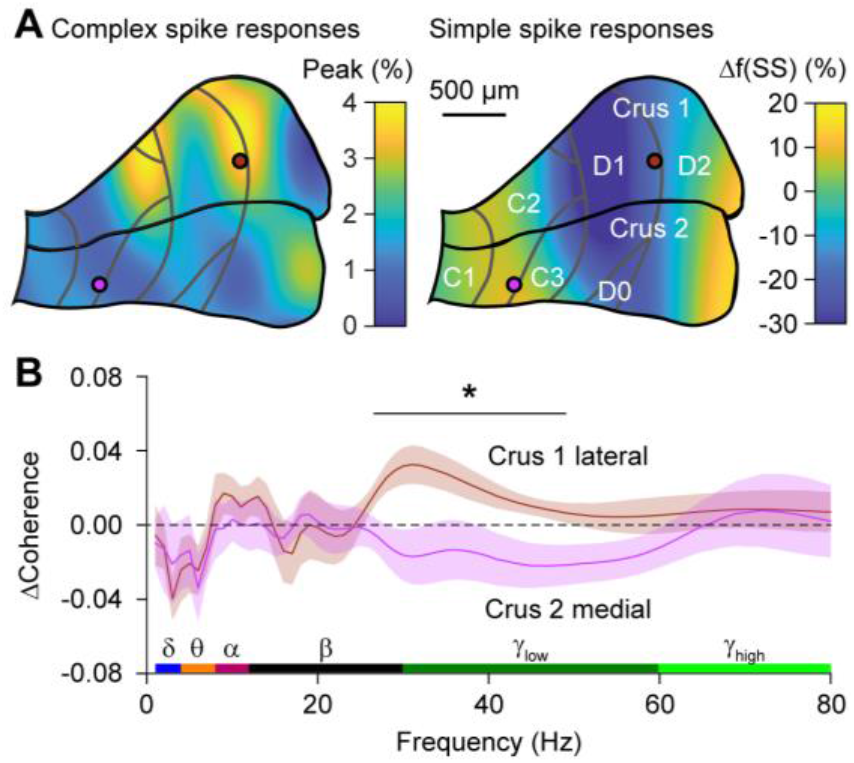
Regional heterogeneity in cerebello-cerebral communication. **A** Air puff whisker pad stimulation results in bidirectional modulation of Purkinje cell simple spike firing. Heat map illustrates the distribution of the maximal modulation within 80 ms of stimulation, showing a difference between medial and lateral zones, modified with permission from (Romano et al., 2018). Note that whisker stimulation can either increase or decrease the simple spike rate. The grey lines indicate the tentative borders between the cerebellar zones. The two colored circles indicate the approximate positions of the 105 μm diameter optic fibers. **B** Compared to air puff stimulation in the absence of optogenetic stimulation, stimulation of Purkinje cells in the medial part of crus 2 and those in the lateral part of crus 1 had opposing effects specifically on sensory-induced gamma band coherence. See also Figs. S8-11.

### Dissecting the impact of neural pathways using a laminar model

We recapitulated the experimental findings by adapting a large-scale computational model of the laminar cortex and subcortical structures (Mejias et al., 2016) to the anatomical pathways relevant for whisker and Purkinje cell stimulation (Fig. 5A). Increasing the intensity of Purkinje cell stimulation had differential effects on different regions, reflecting the contributions of excitatory and inhibitory connections between them (Fig. 5B). Stimulating the trigeminal nucleus, simulating whisker input, induced increased coherence in the theta and lower gamma range, the latter being inhibited by simultaneous Purkinje cell stimulation (Fig. 5C), mimicking the experimental data (see Fig. 2). Granger causality analysis of the modeled data revealed that the sensory-induced gamma band coherence was approximately symmetrical between wS1 and wM1, while the theta band coherence caused by Purkinje cell activity was largely inflicted upon wS1 by wM1 (Fig. 5D). The balance between S1 and M1 in causing sensory-induced gamma band coherence proved to be particularly dependent on the reciprocal connectivity between the superficial layers of S1 and M1 (Fig. S12). Moreover, our model also confirmed that Purkinje cell stimulation is responsible for the enhancement of theta coherence between cortical areas. Given the prominent similarity of the modeled and experimental datasets under various conditions, we next looked at the potential relevance of the different thalamic hubs that were not directly tested in the experiments. These modeling data suggest that the ventroposterior medial nucleus (VPM) and ventrolateral nucleus of the thalamus (VL) had a stronger impact than the medial posterior nucleus (Pom) in mediating the impact of cerebellar activation onto cortical coherence between wS1 and wM1 during sensory stimulation (Fig. S13), which is in line with the distribution of afferents from cerebellum and trigeminal nucleus to the thalamus (Bosman et al., 2011).

**Fig. 5.**
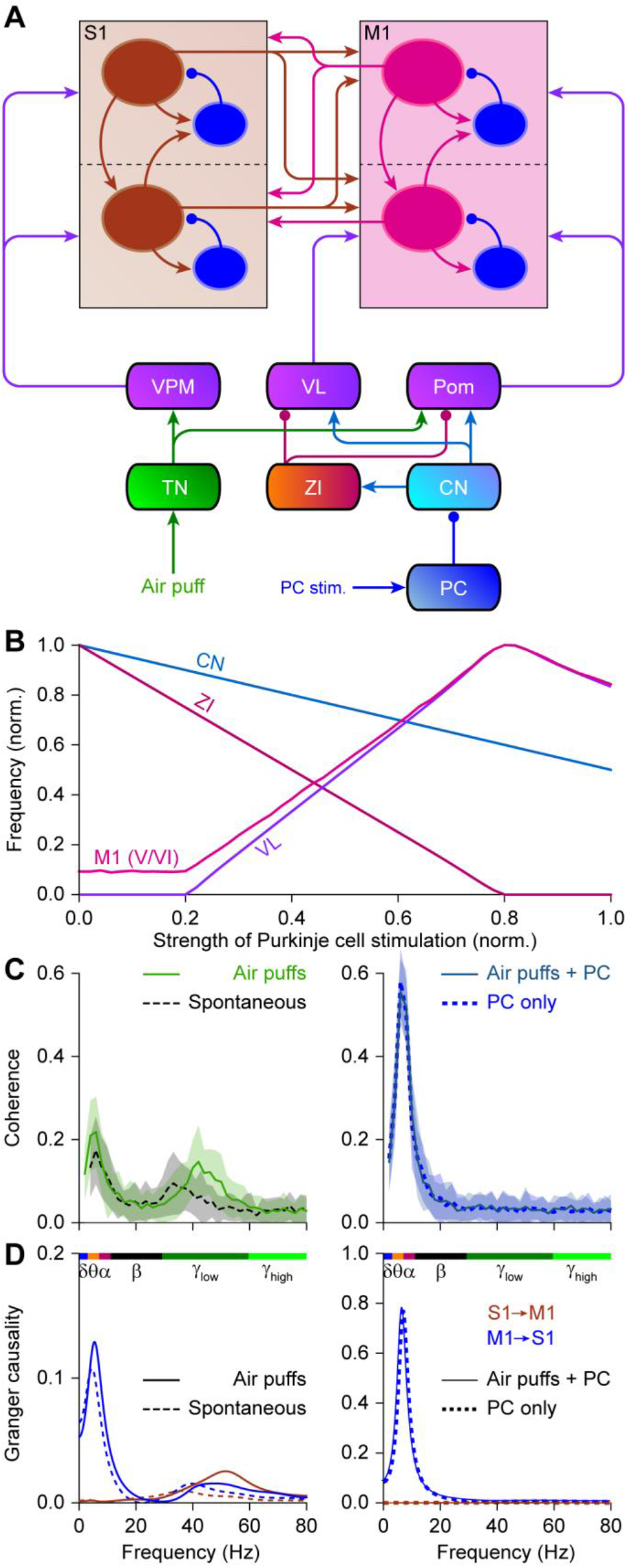
Laminar model. **A** Schematic representation of the connections present in the computational model we used to study cortical coherence *in silico*. CN = cerebellar nuclei, PC = Purkinje cells, Pom = medial posterior nucleus of the thalamus, TN = sensory trigeminal nuclei, VL = ventrolateral nucleus of the thalamus, VPM = ventroposterior medial nucleus of the thalamus, ZI = zona incerta. **B** Impact of Purkinje cell stimulation on the firing rates of four different areas. **C** Coherence between S1 and M1 during stimulation of the trigeminal nuclei (simulating air puffs), the stimulation of Purkinje cells, and the combination of both stimuli. Trigeminal stimulation increased coherence in the gamma band, while Purkinje cell stimulation promoted theta band coherence. Adding Purkinje cell stimulation to trigeminal stimulation cancelled the increased gamma band coherence. Lines represent means and shaded areas sd. **D** Granger causality analysis revealing a largely bidirectional flow between S1 and M1 during gamma band coherence. Lower frequencies predominantly originated from M1. See also Figs. S12 and S13.

## Discussion

Tactile inspection of the world requires acute motor control with fast integration of sensory feedback. This is how one adapts grasp force to keep a slipping cup, or how a tennis player adapts his stroke to side wind. To study fast sensorimotor integration in mammals, exploration by whisker touch has become a popular model system (Bosman et al., 2011; Hartmann, 2011; Prescott et al., 2011). In line with the behavioral relevance to mice, neural control of the facial whiskers is complex, involving synergistic control of cerebellum and neocortex (O’Connor et al., 2002). Here, we found for the first time that transient disruptions of cerebellar output, induced by optogenetic stimulation of Purkinje cells, could disinhibit sensory LFP responses to whisker stimulation in wS1 and wM1 and differentially modulate sensory-induced wS1-wM1 theta and gamma band oscillations. The impact of Purkinje cell stimulation on the coherence in the gamma, but not theta, range depended on the acute behavior as well as the precise location in the cerebellar cortex.

The impact of Purkinje cell stimulation on the individual LFP responses in wS1 and wM1 can be readily explained by the anatomical pathways involved (Bosman et al., 2011). Whisker sensory information is rapidly relayed to the granular layer of wS1 via lemniscal and extralemniscal pathways passing by the thalamic VPM nucleus (Teune et al., 2000; Yu et al., 2006; Urbain and Deschênes, 2007; Furuta et al., 2010; Bosman et al., 2011; Mao et al., 2011; Schäfer and Hoebeek, 2018). Optogenetic silencing of cerebellar output did not affect the initial excitation in the wS1 granular layer, but it did promote the subsequent spread to the subgranular layers (Fig. S4). These effects may be due to the prominent projection of the cerebellar nuclei onto the zona incerta (Teune et al., 2000; Bosman et al., 2011). Indeed, the inhibitory activity of the zona incerta has been implicated in subcortical suppression of whisker sensory wM1.The impact of Purkinje cell stimulation on coherence between wS1 and wM1 showed that cerebellar output enhances and reduces sensory-induced coherence in the theta and gamma bands, respectively. However, the finding that only the impact in the gamma range depended on ongoing behavior and on the precise location of stimulation in the cerebellar cortex raises the possibility that the cerebellum exerts its functional effects in the cortex mainly through a more high-frequency mode of operation. Moreover, these findings also suggest that the cerebellum may better control motor behavior by temporarily downgrading the coherence with sensory relevant signals rather than enhancing them. These implications agree with the high-frequency mode of simple spike activity and modulation that take place during the preparation and execution of motor coordination (Brown and Raman, 2018; Gao et al., 2018; Romano et al., 2018; Chabrol et al., 2019). Indeed, we found that the suppressive impact of Purkinje cell stimulation on gamma band oscillations was greater during larger input during self-motion (Urbain and Deschênes, 2007; Furuta et al., 2010; Bosman et al., 2011; Schäfer and Hoebeek, 2018), which highlights its role as an important intermediate between cerebellum and neocortex. The zona incerta sends GABAergic projections in particular to Pom, which receives direct inputs from the trigeminal nuclei and cerebellum, next to its inputs from wS1 (Bosman et al., 2011; Schäfer and Hoebeek, 2018) and VL (Barthó et al., 2002). Thus, during whisker motion, the cerebellum could, in conjunction with wM1 (Urbain and Deschênes, 2007), activate the zona incerta that in turn suppresses thalamic activity. In wS1, Pom terminals can be found mainly in layers I and V (Zhang and Bruno, 2019), in line with our finding of cerebellar disinhibition of the subgranular layers. In wM1 too, Purkinje cell stimulation resulted in a disinhibition of the sensory response, which again is compatible with the projections from Pom. Moreover, in all cases, a brief delay between sensory and Purkinje cell stimulation reduced the cerebellar impact substantially, which follows our experimental finding that a fast input from the cerebellum appears to be essential for the normal responses in wS1 and movements and this impact could be specifically linked to the Purkinje cells in medial crus 2. Finally, our data also align well with the differential frequencies of coherences that are implicated during the different stages of motor planning and execution (Popa et al., 2013; Arce-McShane et al., 2016).

The impact of cerebellar activation on individual LFP signals in wS1 or wM1 as well as that on the coherence between these signals could be well replicated by our modeling work. We built our computational model on cerebellar modulation of cortical interactions by expanding our existing model on cortico-cortical and thalamo-cortical interactions (Mejias et al., 2016; Jaramillo et al., 2019), The model, the connectivity of which is constrained by realistic anatomical routes, suggests that Purkinje cell activity triggers a disinhibitory effect via the zona incerta, which in turn mediates both the suppression of gamma coherence and the enhancement of theta coherence between S1 and M1. In agreement with the experimental data, the model revealed that sensory-induced gamma band coherence involved mainly signals originating from S1, but with a substantial contribution of M1. The impact of M1 on gamma band coherence depended on the extent of reciprocal connectivity of S1 and M1, rather than on thalamic activity (Figs. S12). Changing the connectivity of the Pom could alter the coherence at lower frequencies, with the stronger connectivity being more in line with the experimental data, but had little impact on the directionality of coherence (Fig. S13). Given the similarities between the modeling and experimental outcomes, even explaining counterintuitive findings, the current model may well provide detailed and valid predictions as to how the cerebellum may influence the different layers and areas of the cerebral cortex under a wider and richer variety of physiological behaviors.

## Materials and Methods

### Animals

Experiments were performed on heterozygous transgenic mice expressing the light-sensitive cation channel channelrhodopsin-2 under the Purkinje cell-specific *Pcp2* promoter (Tg(Pcp2-cre)2MPin;Gt(ROSA)26Sor^tm27.1(CAG-COP4*H134R/tdTomato)Hze^) on a C57BL6/J background (Witter et al., 2013). We used 14 males and 12 females aged between 10 and 34 weeks. The mice were kept in a vivarium with controlled temperature and humidity and a 12/12 h light/dark cycle. The animals were group housed until surgery and single housed afterwards. A project license was obtained prior to the start of the experiments from the national authority (Centrale Commissie Dierproeven, The Hague, The Netherlands; license no. AVD101002015273) as required by Dutch law and all experiments were performed according to institutional, national and EU guidelines and legislation.

### Surgery

The mice received a pedestal to allow head-fixation in the recording setup as well as one to three craniotomies to grant access to the brain. All surgical procedures were performed under anesthesia and the mice were given pain-killers before and after surgery. The animals were given three days of recovery after the surgery before they were habituated to the setup on at least three consecutive days with increasing habituation times (from approx. 10 min the first session to approx. 2 h the last session). Further details can be found in the Supplementary Methods.

### Electrophysiology

All recordings were made in awake, head restrained mice. Single unit activity of putative cerebellar nuclei neurons was measured using extracellular quartz-coated platinum-tungsten fiber electrodes (R = 2-5 MΩ; 80 μm outer diameter; Thomas Recording, Giessen, Germany) placed in a rectangular matrix (Thomas Recording) with an inter-electrode distance of 305 μm. LFP recordings were made in wS1 and wM1 using 16 channel, single shaft silicon probes with an inter-electrode distance of 100 μm (R = 1.5-2.5 MΩ, A1×16-5mm-100-177-A16, NeuroNexus Technologies, Ann Arbor, MI, USA). Each silicon probe was equipped with its own reference, placed in close proximity to the recording site. The two probes shared the same ground, which was placed either in the agar covering the recording sites or in the agar covering the cerebellar craniotomy. All electrodes were connected to a PZ5 NeuroDigitizer (Tucker-Davis Technologies). The signals were amplified, 1-6,000 Hz filtered, digitized at 24 kHz and stored using a RZ2 multi-channel workstation (Tucker-Davis Technologies). Recorded neurons were classified as putative cerebellar nuclei neurons if they were recorded at a depth of at least 1700 μm from the cerebellar surface and if the recording contained only a single type of action potentials typically showing both negative and positive parts, what differentiated them from Purkinje cell simple spikes. Spike times from single-unit recordings were retrieved off-line using Spiketrain (Neurasmus BV, Rotterdam, The Netherlands). Before any analysis was done on the LFP data, the raw traces were normalized using the z-score function in Matlab (MathWorks, Nattick, MA, USA). The current source density analysis was performed in custom written Matlab routines as detailed in the Supplementary Methods.

### Coherence analysis

The phase coherence analysis was computed using the Fieldtrip toolbox as described in the Supplementary Methods. For this, LFP snippets of 5 second pre- and 5 second post-stimulus were used to calculate the coherence spectrum per trial. If necessary, 50 Hz line noise was removed. Next, the coherence in a 100 ms window after stimulus onset was averaged per frequency. The effect of Purkinje cell activation on the sensory triggered wS1-wM1 coherence was determined by subtracting the averaged air puff-induced coherence from the air puff with photostimulation evoked coherence. The Granger causality analysis was carried out by the Fieldtrip toolbox with the same preprocessing on the LFP data.

### Stimulation

Optogenetic stimulation of the cerebellum occurred contralateral to the neocortical LFP recording sites using 470 nm LED drivers (M4703F, ThorLabs. Newton, NJ, USA) connected to a 4-channel LED driver (DC4104, ThorLabs) and optic fibers with diameters of 400 (Figs. 1–3) or 105 μm (Fig. 4) (ThorLabs). The 400 μm fibers were placed just above the dura of the cerebellum. The 105 μm fibers were adapted for insertion into the rectangular electrode matrix by removing the cladding for ~15 cm and grinding the tip under microscope guidance. Unless stated otherwise, photostimulation was applied as 100 ms pulses with a power of 7.0 mW (400 μm fiber) or 0.2 mW (105 μm fiber). Sensory stimulations consisted of 30 ms air puffs at 1 bar directed at the mystacial macrovibrissae ipsilateral to cerebellar and contralateral to neocortical recording sites, using a MPPI-2 pressure injector (Applied Scientific Instrumentation, Eugene, OR, USA). The nozzle was positioned to minimize stimulation of the eye or ear. Stimuli were presented at 0.25 Hz in pseudorandom order.

### Whisker movement tracking

Whisker movements in awake head-restrained mice were recorded with a high-speed video camera (frame rate 1,000 Hz; A504k camera, Basler, Ahrensburg, Germany), using a custom-made LED panel (λ = 640 nm) as back-light. All whiskers were kept intact. We tracked the whiskers as described previously (Rahmati et al., 2014; Romano et al., 2018). For this study, the whisker position was defined as the average angle of all trackable whiskers. See the Supplementary Methods for more details.

### Computational model

The computational model used is based on the one developed in (Mejias et al., 2016), with (i) minimal variations in the cortical parameters based on the observed anatomical and physiological properties of wS1 and wM1 in mice and (ii) the addition of trigeminal nucleus, thalamic nuclei and cerebellar areas to the network.

#### Neocortex

Each cortical area is constituted by two cortical layers (or more generally, laminar modules) which describe the dynamics of superficial and deep layers, respectively. A laminar module contains one excitatory and one inhibitory population, and the dynamics of their respective firing rates *r_E_*(*t*) and *r_I_*(*t*) are described by the following equations:

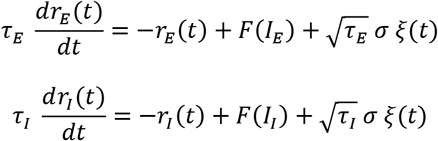

Here, *τ_E_*, *τ_I_* denote the time scales for the excitatory and inhibitory populations respectively, and *ξ_E_*(*t*), *ξ_I_*(*t*) are Gaussian white noise terms of zero mean and standard deviation *σ*. For superficial layers, we choose *τ_E_* = 6 ms, *τ_I_* = 15 *ms* and *σ* = 0.3, which leads to a noisy oscillatory dynamics in the gamma range, and for deep layers we choose *τ_E_* = 48 *ms*, *τ_I_* = 120 *ms* and *σ* = 0.45, which leads to noisy oscillations in the theta and low alpha range. Note that the relatively high values for the time constants in deep layers are thought to reflect other slow biophysical factors not explicitly included in the model, such as the dynamics of NMDA receptors.

The function *F*(*x*) = *x*/(1 − exp(−*x*)) is the input-output transfer function of each population, which transforms the incoming input currents into their corresponding cell-averaged firing rates. The argument of the transfer function is the incoming current for each population, which involves a background term, a local term and a long-range term. The background term is a default constant current only received by excitatory neurons in S1 and M1, and it is *I_bg_ = 4* for superficial excitatory neurons and *I_bg_ = 1* for deep excitatory neurons. The local term involves the input coming from neurons within the area, and it is given by

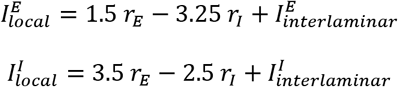

Here, the numbers denote the strengths of the synaptic projections considered. The interlaminar terms are contributions from a different layer than the one the population is in. The only interlaminar projections are from superficial excitatory to deep excitatory neurons, with synaptic strength 1, and from deep excitatory to superficial inhibitory neurons, with synaptic strength 0.75 (Kros et al., 2015).

Finally, the long-range term includes currents coming from other neocortical or subcortical areas. These currents follow the general form *J_ab_r_b_*, (with *J_ab_* being the synaptic strength from area ‘b’ to area ‘a’) and therefore we will specify only the synaptic coupling strengths to characterize them.

Following anatomical evidence (Bosman et al., 2011), we consider excitatory projections from superficial S1 neurons to both superficial (strength 0.52) and deep (0.25) excitatory M1 neurons, and from deep S1 neurons to superficial (0.25) and deep (0.75) excitatory M1 neurons. In the opposite direction, we consider excitatory projections from superficial M1 neurons to both superficial (0.5) and deep (1) S1 excitatory neurons, and from deep M1 neurons to deep (1) S1 excitatory neurons.

The dynamics of the firing rate of the trigeminal nucleus (TN), the thalamic nuclei (VPM, VL and Pom) and cerebellar populations (PC, CN and ZI) are each described by equations of the type

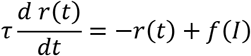

Here, *τ* = 10 *ms* is the characteristic time constant and the transfer function is a threshold-linear function (i.e. *f*(*x*) = *Ax*, with A being the gain of the population, for x>0, and *f*(*x*) = 0 otherwise). The gain parameter A takes the values 3, 10, 1, 1, 0.5, 5 and 0.2 for areas TN, PC, CN, ZI, VPM, VL and Pom, respectively. Air puffs are modeled as a constant input (max I=10) to TN, while optogenetic stimulation to PC is modeled as a constant input (I=1). Cerebellar areas CN and ZI receive inhibitory projections (both of strength 1) from PC and CN respectively. In addition, PC and CN received excitatory background currents of 0.1 and 21 respectively, and ZI receives an inhibitory background current of 12 (which can be also interpreted as a high firing threshold). Thalamic nuclei VPM received an excitatory projection (strength 1) from TN, VL receives projections from CN (strength 1) and ZI (strength −3), and Pom receives projections from TN (1), CN (0.2) and ZI (−0.5). Projections from VPM reach superficial excitatory (strength 0.66), deep excitatory (0.13) and inhibitory (0.2) populations of S1. Regarding M1, it receives projections from Pom to all its excitatory (0.33) and inhibitory (0.5) populations, and deep excitatory M1 neurons also receive a projection (0.6) from VL. When projections from Pom to S1 are considered (see Fig. S13), they target both excitatory (0.2) and inhibitory (0.15) populations in S1.

To mimic the depth of the recording electrodes for wS1 and wM1 in experiments, we estimate the LFP signal in the model by a weighted average of the excitatory superficial and deep layers, with a superficial:deep ratio of 1:9 for wS1 (i.e. deep layers) and 4:6 for wM1 (as it targets more superficial layers but it would still pick up signals from apical dendrites’ layer V neurons).

### Experimental design and statistical analysis

We considered *p* ≤ 0.05 as significant unless Benjamini-Hochberg correction for multiple comparisons was applied (see Table S1). Two-tailed testing was used for all statistical analyses. *N* indicates the number of mice; *n* indicates the number of stimuli/recordings. Further details are in the Supplementary Methods.

## Acknowledgments

Financial support was provided by the Netherlands Organization for Scientific Research (NWO-ALW; CIDZ), the Dutch Organization for Medical Sciences (ZonMW; CIDZ), Life Sciences (CIDZ), ERC-adv and ERC-POC (CIDZ), as well as Crossover NWO-LSH INTENSE and Medical NeuroDelta. SH was supported by funding from the Okinawa Institute of Science and Technology Graduate University.

## References

Ahrens KF, Kleinfeld D (2004) Current flow in vibrissa motor cortex can phase-lock with exploratory rhythmic whisking in rat. J Neurophysiol 92:1700–1707.

Arce-McShane FI, Ross CF, Takahashi K, Sessle BJ, Hatsopoulos NG (2016) Primary motor and sensory cortical areas communicate via spatiotemporally coordinated networks at multiple frequencies. Proc Natl Acad Sci U S A 113:5083–5088.

Barthó P, Freund TF, Acsády L (2002) Selective GABAergic innervation of thalamic nuclei from zona incerta. Eur J Neurosci 16:999–1014.

Bosman LWJ, Koekkoek SKE, Shapiro J, Rijken BFM, Zandstra F, van der Ende B, Owens CB, Potters JW, de Gruijl JR, Ruigrok TJH, De Zeeuw CI (2010) Encoding of whisker input by cerebellar Purkinje cells. J Physiol 588:3757–3783.

Bosman LWJ, Houweling AR, Owens CB, Tanke N, Shevchouk OT, Rahmati N, Teunissen WHT, Ju C, Gong W, Koekkoek SKE, De Zeeuw CI (2011) Anatomical pathways involved in generating and sensing rhythmic whisker movements. Front Integr Neurosci 5:53.

Bressler SL, Coppola R, Nakamura R (1993) Episodic multiregional cortical coherence at multiple frequencies during visual task performance. Nature 366:153–156.

Brown ST, Raman IM (2018) Sensorimotor integration and amplification of reflexive whisking by well-timed spiking in the cerebellar corticonuclear circuit. Neuron 99:564–575.

Buzsáki G, Wang XJ (2012) Mechanisms of gamma oscillations. Annu Rev Neurosci 35:203–225.

Cardin JA, Carlén M, Meletis K, Knoblich U, Zhang F, Deisseroth K, Tsai LH, Moore CI (2009) Driving fast-spiking cells induces gamma rhythm and controls sensory responses. Nature 459:663–667.

Chabrol FP, Blot A, Mrsic-Flogel TD (2019) Cerebellar contribution to preparatory activity in motor neocortex. Neuron 103:506–519 e504.

De Kock CPJ, Bruno RM, Spors H, Sakmann B (2007) Layer and cell type specific suprathreshold stimulus representation in primary somatosensory cortex. J Physiol 581:139.

De Zeeuw CI, Hoebeek FE, Bosman LWJ, Schonewille M, Witter L, Koekkoek SK (2011) Spatiotemporal firing patterns in the cerebellum. Nat Rev Neurosci 12:327–344.

Engel AK, König P, Kreiter AK, Schillen TB, Singer W (1992) Temporal coding in the visual cortex: new vistas on integration in the nervous system. Trends Neurosci 15:218–226.

Ferezou I, Haiss F, Gentet LJ, Aronoff R, Weber B, Petersen CCH (2007) Spatiotemporal dynamics of cortical sensorimotor integration in behaving mice. Neuron 56:907–923.

Fries P (2015) Rhythms for Cognition: Communication through Coherence. Neuron 88:220–235.

Furuta T, Urbain N, Kaneko T, Deschênes M (2010) Corticofugal control of vibrissa-sensitive neurons in the interpolaris nucleus of the trigeminal complex. J Neurosci 30:1832–1838.

Gao Z, Davis C, Thomas AM, Economo MN, Abrego AM, Svoboda K, De Zeeuw CI, Li N (2018) A cortico-cerebellar loop for motor planning. Nature 563:113–116.

Hartmann MJZ (2011) A night in the life of a rat: vibrissal mechanics and tactile exploration. Annals of the New York Academy of Sciences 1225:110–118.

Jaramillo J, Mejias JF, Wang XJ (2019) Engagement of pulvino-cortical feedforward and feedback pathways in cognitive computations. Neuron 101:321–3336 e329.

Jensen O, Spaak E, Zumer JM (2014) Human brain oscillations: From physiological mechanisms to analysis and cognition. In: Magnetoencephalography: From Signals to Dynamic Cortical Networks (Supek S, Aine CJ, eds), pp 359–403. Berlin, Heidelberg: Springer Berlin Heidelberg.

Kleinfeld D, Sachdev RNS, Merchant LM, Jarvis MR, Ebner FF (2002) Adaptive filtering of vibrissa input in motor cortex of rat. Neuron 34:1021–1034.

Kros L, Eelkman Rooda OHJ, Spanke JK, Alva P, van Dongen MN, Karapatis A, Tolner EA, Strydis C, Davey N, Winkelman BHJ, Negrello M, Serdijn WA, Steuber V, van den Maagdenberg AMJM, De Zeeuw CI, Hoebeek FE (2015) Cerebellar output controls generalized spike-and-wave discharge occurrence. Ann Neurol 77:1027–1049.

Mao T, Kusefoglu D, Hooks BM, Huber D, Petreanu L, Svoboda K (2011) Long-range neuronal circuits underlying the interaction between sensory and motor cortex. Neuron 72:111–123.

Mejias JF, Murray JD, Kennedy H, Wang XJ (2016) Feedforward and feedback frequency-dependent interactions in a large-scale laminar network of the primate cortex. Sci Adv 2:e1601335.

O’Connor SM, Berg RW, Kleinfeld D (2002) Coherent electrical activity between vibrissa sensory areas of cerebellum and neocortex is enhanced during free whisking. J Neurophysiol 87:2137–2148.

Pedroarena C, Llinás R (1997) Dendritic calcium conductances generate high-frequency oscillation in thalamocortical neurons. Proc Natl Acad Sci U S A 94:724–728.

Pettersen KH, Hagen E, Einevoll GT (2008) Estimation of population firing rates and current source densities from laminar electrode recordings. J Comput Neurosci 24:291–313.

Popa D, Spolidoro M, Proville RD, Guyon N, Belliveau L, Léna C (2013) Functional role of the cerebellum in gamma-band synchronization of the sensory and motor cortices. J Neurosci 33:6552–6556.

Prescott TJ, Diamond ME, Wing AM (2011) Active touch sensing. Philos Trans R Soc Lond B Biol Sci 366:2989–2995.

Proske U, Gandevia SC (2009) The kinaesthetic senses. J Physiol 587:4139–4146.

Proville RD, Spolidoro M, Guyon N, Dugué GP, Selimi F, Isope P, Popa D, Léna C (2014) Cerebellum involvement in cortical sensorimotor circuits for the control of voluntary movements. Nat Neurosci 17:1233–1239.

Rahmati N, Owens CB, Bosman LWJ, Spanke JK, Lindeman S, Gong W, Potters JW, Romano V, Voges K, Moscato L, Koekkoek SKE, Negrello M, De Zeeuw CI (2014) Cerebellar potentiation and learning a whisker-based object localization task with a time response window. J Neurosci 34:1949–1962.

Romano V, De Propris L, Bosman LWJ, Warnaar P, ten Brinke MM, Lindeman S, Ju C, Velauthapillai A, Spanke JK, Middendorp Guerra E, Hoogland TM, Negrello M, D’Angelo E, De Zeeuw CI (2018) Potentiation of cerebellar Purkinje cells facilitates whisker reflex adaptation through increased simple spike activity. eLife 7:e38852.

Saalmann YB, Pinsk MA, Wang L, Li X, Kastner S (2012) The pulvinar regulates information transmission between cortical areas based on attention demands. Science 337:753–756.

Schäfer CB, Hoebeek FE (2018) Convergence of primary sensory cortex and cerebellar nuclei pathways in the whisker system. Neuroscience 368:229–239.

Shin H, Moore CI (2019) Persistent gamma spiking in SI nonsensory fast spiking cells predicts perceptual success. Neuron.

Song W, Francis JT (2015) Gating of tactile information through gamma band during passive arm movement in awake primates. Frontiers in neural circuits 9:64.

Teune TM, van der Burg J, van der Moer J, Voogd J, Ruigrok TJ (2000) Topography of cerebellar nuclear projections to the brain stem in the rat. Progress in brain research 124:141–172.

Urbain N, Deschênes M (2007) A new thalamic pathway of vibrissal information modulated by the motor cortex. J Neurosci 27:12407–12412.

van Kerkoerle T, Self MW, Dagnino B, Gariel-Mathis MA, Poort J, van der Togt C, Roelfsema PR (2014) Alpha and gamma oscillations characterize feedback and feedforward processing in monkey visual cortex. Proc Natl Acad Sci U S A 111:14332–14341.

Vecchio F, Caliandro P, Reale G, Miraglia F, Piludu F, Masi G, Iacovelli C, Simbolotti C, Padua L, Leone E, Alù F, Colosimo C, Rossini PM (2019) Acute cerebellar stroke and middle cerebral artery stroke exert distinctive modifications on functional cortical connectivity: A comparative study via EEG graph theory. Clinical Neurophysiology 130:997–1007.

Veit J, Hakim R, Jadi MP, Sejnowski TJ, Adesnik H (2017) Cortical gamma band synchronization through somatostatin interneurons. Nat Neurosci 20:951–959.

Witter L, Canto CB, Hoogland TM, de Gruijl JR, De Zeeuw CI (2013) Strength and timing of motor responses mediated by rebound firing in the cerebellar nuclei after Purkinje cell activation. Frontiers in neural circuits 7:133.

Womelsdorf T, Fries P (2006) Neuronal coherence during selective attentional processing and sensorymotor integration. J Physiol Paris 100:182–193.

Yu C, Derdikman D, Haidarliu S, Ahissar E (2006) Parallel thalamic pathways for whisking and touch signals in the rat. PLoS Biol 4:e124.

Zhang W, Bruno RM (2019) High-order thalamic inputs to primary somatosensory cortex are stronger and longer lasting than cortical inputs. Elife 8:e44158.

