## Supplementary information for "Cerebellar Purkinje cells can differentially modulate coherence between sensory and motor cortex depending on region and behavior"

### Coherence bands between primary sensory and motor cortex can be differentially modulated by cerebellar Purkinje cells in a region-specific and behavior-dependent manner

Sander Lindeman, Lieke Kros, Sungho Hong, Jorge F. Mejias, Vincenzo Romano, Mario Negrello, Laurens W.J. Bosman and Chris I. De Zeeuw

Laurens Bosman

Mario Negrello


**Supplementary methods**

**Surgery.** All surgical procedures were performed under anesthesia (2-5% isoflurane in 1 l/min oxygen) in combination with treatment of surgical pain giving 5 mg/kg carprofen (“Rimadyl”, Pfizer, New York, NY, USA), 1 µg bupivacaine (Actavis, Parsippany-Troy Hills, NJ, USA) and 50 µg/kg buprenorphine ("Temgesic”, Indivior, Richmond, VA, USA). In addition, the mice received 1 µg lidocaine (Braun, Meisingen, Germany) subcutaneously at the surgical sites prior to the start of the surgery. The body temperature was maintained at 37 °C by a feed-back controlled heating pad.

For the placement of a pedestal, a part of the skin was removed and the skull was cleaned and treated with phosphoric acid to ensure all membranes were removed. Next, the exposed skull was treated with Optibond adhesive (Kerr Dental, Orange, CA, USA) and the mice received a magnetic pedestal that was placed on the skull between the eyes and secured with Charisma (Kerr Dental). Next, up to three craniotomies were performed allowing access to the whisker part of the left primary somatosensory (wS1, relative to bregma: 3.5 mm mediolateral and -1.5 mm anteroposterior) and motor cortex (wM1, relative to bregma: 1.5 mm mediolateral and 1.0 mm anteroposterior) and the right cerebellar hemisphere, each surrounded by a recording chamber made out of Charisma. The exposed dura was covered with tetracycline-containing ointment (Terra Cortril; Pfizer, New York, NY, USA) and the recording chambers were sealed with a silicon polymer (Kwik-Cast, WPI, Sarasota, FL, USA) and covered with bone wax (Ethicon, Somerville, NJ, USA). The animals were given three days of recovery after the surgery before they were habituated to the setup on at least three consecutive days with increasing habituation times (from approx. 10 min the first session to approx. 2 h the last session).

**Electrophysiology.** Local field potentials (LFP) were recorded in wS1 and wM1 using linear silicon probes. Each silicon probe was equipped with its own reference, placed in close proximity to the recording site. The two probes shared the same ground, which was placed either in the agar covering the recording sites or in the agar covering the cerebellar craniotomy. The platinum-tungsten electrodes as well as the silicon probes were connected to a PZ5 NeuroDigitizer (Tucker-Davis Technologies (TDT), Alachua, FL, USA). The signals were amplified, 1-6,000 Hz filtered, digitized at 24 kHz and stored using a RZ2 multi-channel workstation (TDT).

Before any analysis was done on the LFP data, the raw traces were normalized using the z-score function in MATLAB (MathWorks, Nattick, MA, USA). The current source density analysis was performed in custom written MATLAB routines using the Kernel Source Density Method as described in (1); see https://github.molgen.mpg.de/MPIBR-coattia/MatlabMain/tree/master/behaviorAnalysis/code/functions/kCSDv1.

In the cerebellum, recorded neurons were classified as originating from putative cerebellar nuclei neurons if they were recorded at a depth of at least 1700 µm from the cerebellar surface and if the recording contained only a single type of action potentials, what differentiated them from Purkinje cell recordings. Spike times were retrieved off-line using SpikeTrain (Neurasmus BV, Rotterdam, The Netherlands). After automated spike detection and sorting, all traces were inspected manually and improper event classification was corrected.

**Coherence analysis.** The phase coherence analysis was computed using the Fieldtrip toolbox (2). For this, LFP snippets of 5 second pre- and 5 second post-stimulus were used to calculate the coherence spectrum per trial. If necessary, line noise at 50 Hz was removed first from the waveforms by fitting a PSD around the time of the peaks of the power spectrum and then filtering the signal with the inversed square root of this function. Next, the coherence in a frequency-dependent window (2 * 1/frequency) after stimulus onset was averaged per frequency to perform the further analysis on. The effect of optogenetic Purkinje cell activation on the sensory triggered wS1-wM1 coherence was determined by subtracting the averaged air puff induced coherence from the air puff with optogenetically evoked coherence. To test for differences between the conditions, the difference of coherence test was used, as described by Amjad et al. (3). In short, the Fisher transform ($\tanh^{-1}$) was applied on the coherence and this was compared to a $\chi^2$-distribution with k - 1 degrees of freedom, where k is the number of conditions that were tested (in all cases k = 2). The 95% confidence limit was then determined using $\chi^2_{(0.05;1)} = 3.84$. The significant frequencies are indicated in the difference of coherence figures using lines and asterisks.

**Whisking behavior.** Whisker movements were tracked off-line using the BIOTACT Whisker Tracking Tool (BWTT) with the sdGeneric, stShapeSpaceKalman, ppBigExtractionAndFiltering, and wdIgorMeanAngle plugins (http://bwtt.sourceforge.net) (4). Briefly, we first determined the position of the snout in each frame semi-automatically by fitting a template to the snout. After masking the snout and subtracting the unmoved background from each frame, the whiskers were traced in a radial approach. The algorithm detected edges in the frame in consecutive concentric snout-shaped masks around the actual snout mask. Ultimately, we detected the start and end nodes of the fitted line segments, and calculated the angles of the whiskers from these values. The final BWTT result provided us with the angles of all detected whiskers per video frame. To relate the angles across frames to the tracks, we wrote an algorithm that predicts track values in consecutive frames based on the position and velocity in the angular value as well as the y-position of the last video frames (5, 6). The predicted track values for the next frame were compared with the detected values in the next frame and were assigned according to a minimum deviation approach between them. Finally, the mean angle per frame was calculated from the individual whisker traces.

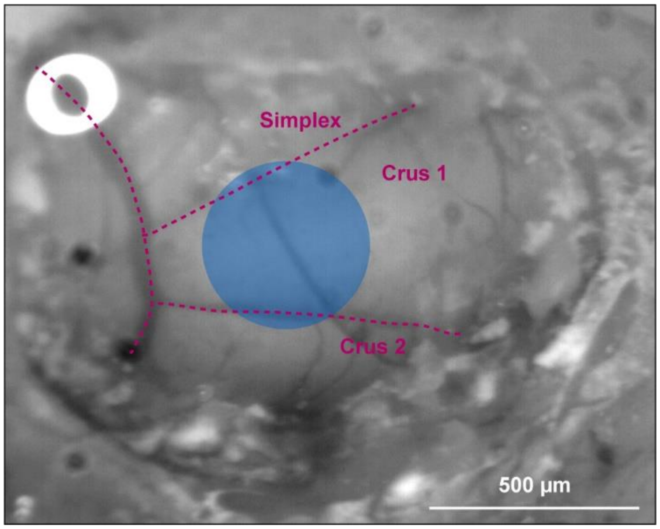


**Fig. S1. Location of optogenetic stimulation.**

Approximate location of a 400 µm diameter optic fiber in the center of crus 1 as seen through the craniotomy.

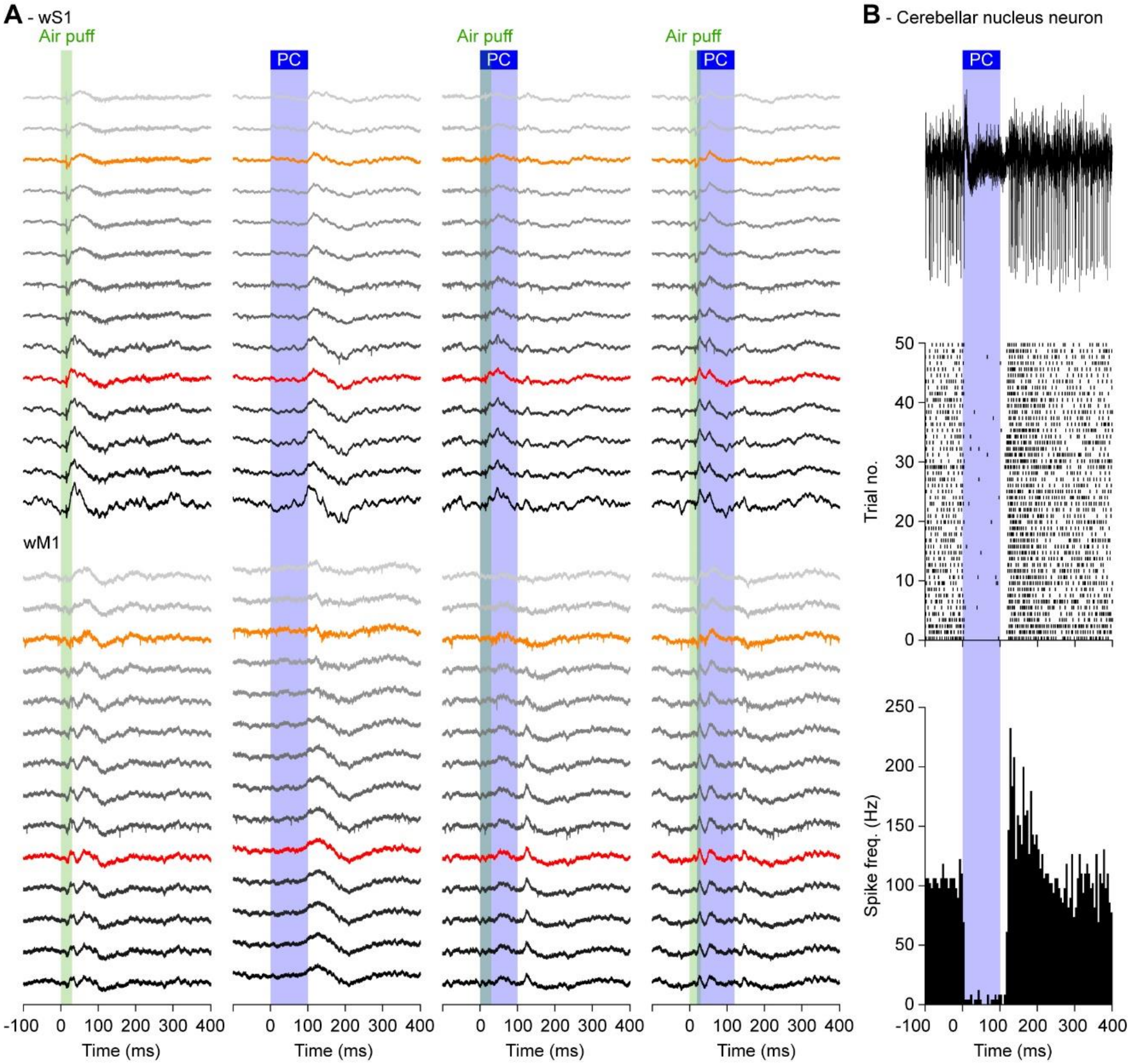


**Fig. S2.** Purkinje cell stimulation reduces the impact of whisker stimulation on wS1 and wM1.

**A** Air puff stimulation of the large facial whiskers evoked sensory responses in contralateral wS1 and wM1, recorded here as deviations in the local field potential (LFP) of a randomly selected single trial (left column). The LFP recordings were made using linear silicon probes with 100 µm inter-electrode distances. The recordings are organized from superficial to deep (color code as in Fig. 1B). Electrodes 3 and 10, that were used for most analyses in this study, are marked with orange and red, respectively. Optogenetic stimulation of Purkinje cells (PC) was done with an optic fiber with a diameter of 400 µm placed on the center of crus 1 (see Fig. S1), leading to a delayed response in both wS1 and wM1 (2nd column). The other columns depict randomly selected trials from the same experiment, showing respectively the combined sensory and optogenetic Purkinje cell stimulation and the latter with a delay of 20 ms before the onset of the Purkinje cell stimulation. **B** Optogenetic Purkinje cell stimulation leads to a pause in firing of an exemplary neuron in the cerebellar nuclei, followed by rebound firing after the end of stimulation. Further analysis of cerebellar nuclear activity is presented in Figs. S8-S10.

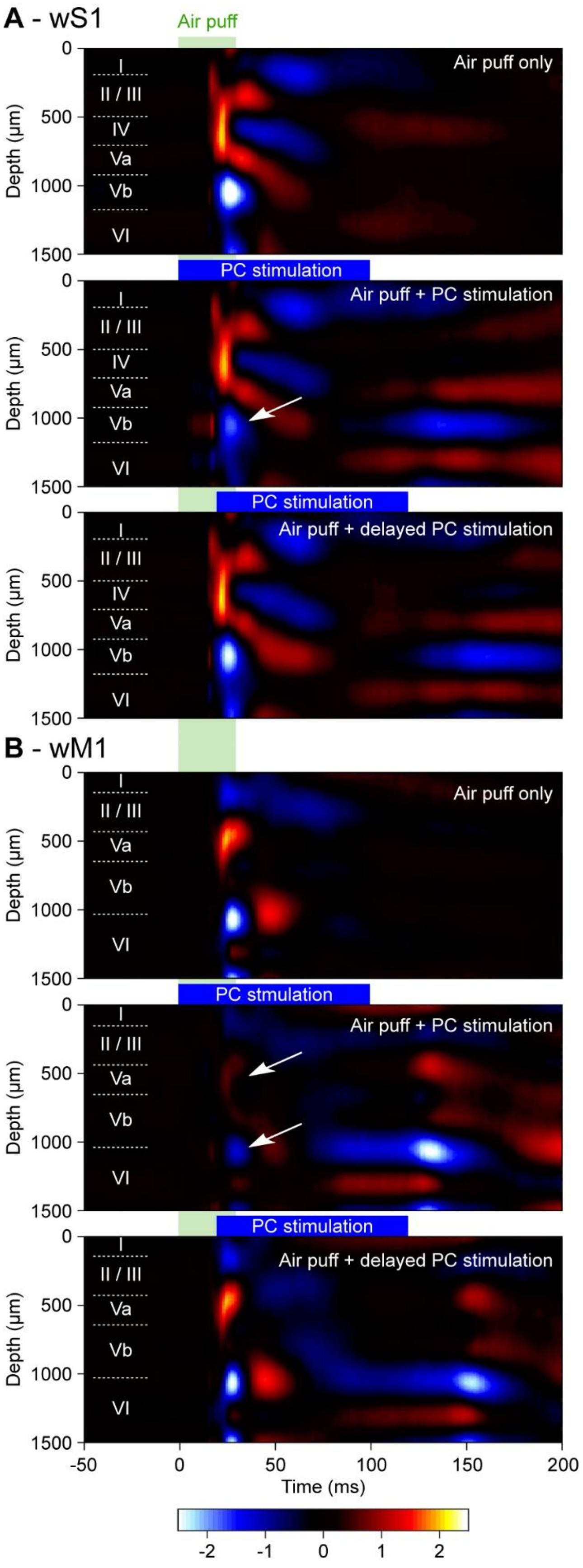


**Fig. S3. Purkinje cell stimulation reduces sensory responses in wS1 and wM1.**

**A** Current source density analysis of the averaged local field potentials across the layers of the whisker area of S1 (see Fig. 1C-H) during whisker air puff stimulation alone or in combination with optogenetic stimulation of Purkinje cells using an optic fiber with a diameter of 400 µm placed on the center of crus 1 (see Fig. S1). Purkinje cell stimulation applied simultaneously with whisker stimulation suppressed predominantly the fast current sinks (blue; white arrow), but a 20 ms delay between air puff and whisker stimulation largely restores the impact of air puff stimulation. **B** The same was true for the whisker area of wM1. The heat maps indicate the averaged values of 8 mice with color scaling in arbitrary units. The layers of wS1 and wM1 are indicated by approximation.

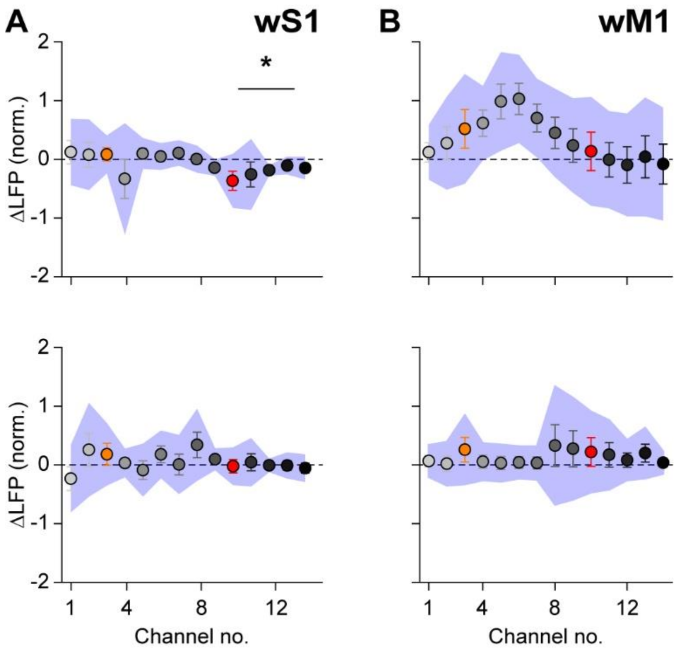


**Fig. S4. Optogenetic Purkinje cell stimulation increases the spread of excitation triggered by whisker stimulation.**

**A** Air puff whisker stimulation triggers excitation in contralateral wS1 and wM1. In wS1, the amplitude of the first negative LFP peak (corresponding to excitatory activity) in the subgranular layers was enhanced ($p$ = 0.012, $\chi^2$ = 2.500, Dunn's post-hoc test after Friedmann's ANOVA, Table S1). This effect was absent upon introducing a delay of 20 ms between whisker and Purkinje cell stimulation (bottom row). **B** The same for wM1. Plotted are the averaged differences in amplitude of the first positive LFP peaks. Error bars indicate SEM and shaded area sd. $n$ = 100 trials each in $N$ = 8 mice.

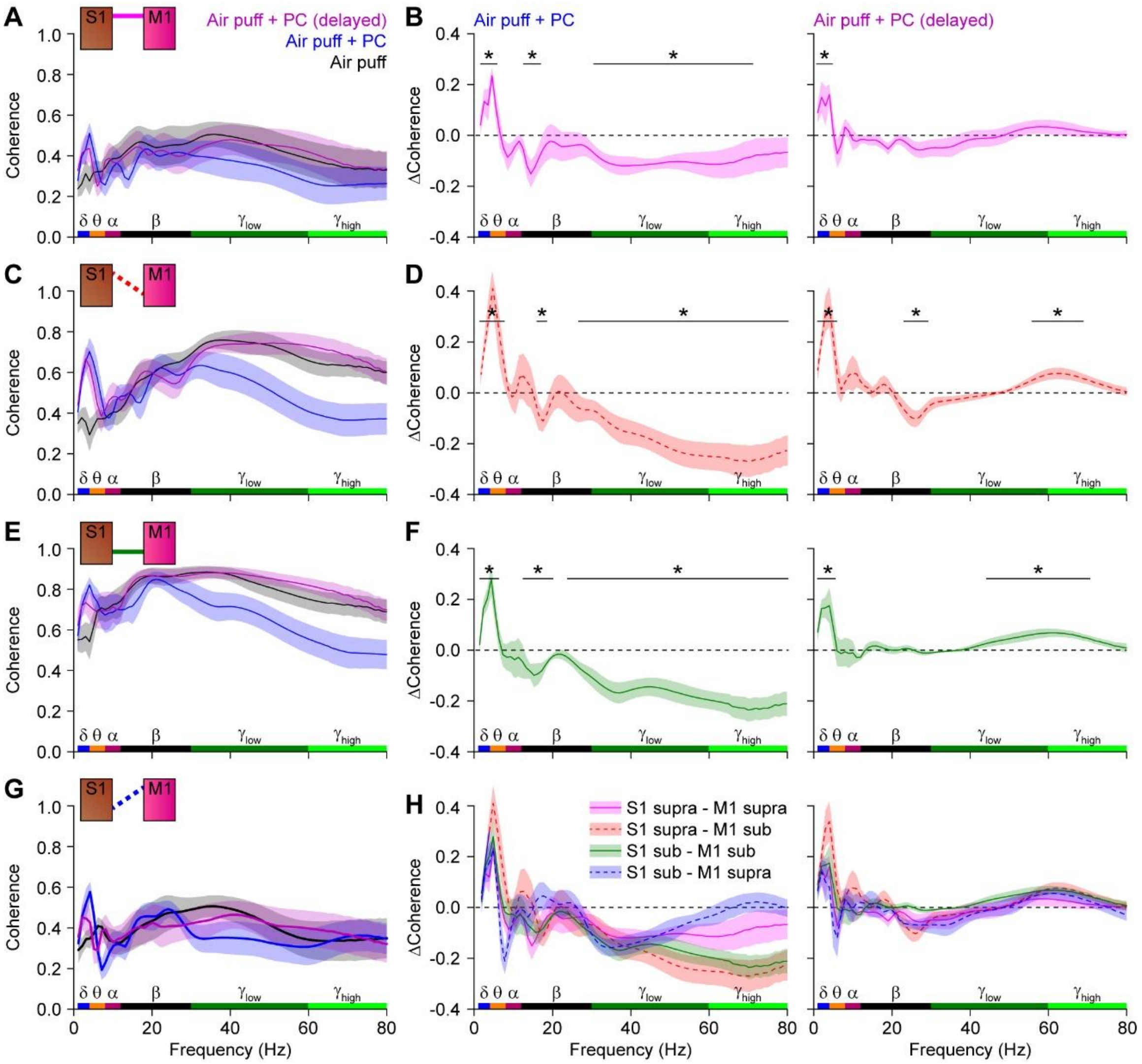


**Fig. S5. Cerebellar Purkinje cell stimulation suppresses sensory-induced gamma band coherence between wS1 and wM1.**

**A** Averaged coherence between the supragranular layers of wS1 and wM1 following air puff stimulation of the contralateral facial whiskers in isolation or in combination with simultaneous or 20 ms delayed optogenetic stimulation of Purkinje cells using an optic fiber with a diameter of 400 µm placed on the center of crus 1 (see Fig. S1). Shaded areas indicate SEM. $n$ = 100 trials per condition each in $N$ = 8 mice. **B** Purkinje cell stimulation suppressed mainly the gamma band coherence induced by air puff sensory stimulation. This effect was largely abolished by introducing a 20 ms delay between the start of the sensory stimulation and that of the Purkinje cells. **C**-**H** The same for the coherence between different layers of wS1 and wM1 as indicated schematically in the upper left corners. Although the details varied to some extent, in all cases Purkinje cell stimulation suppressed sensory-induced gamma band coherence between wS1 and wM1. * $p < 0.05$ ($\chi^2 > 3.84$; difference of coherence test, see Methods).

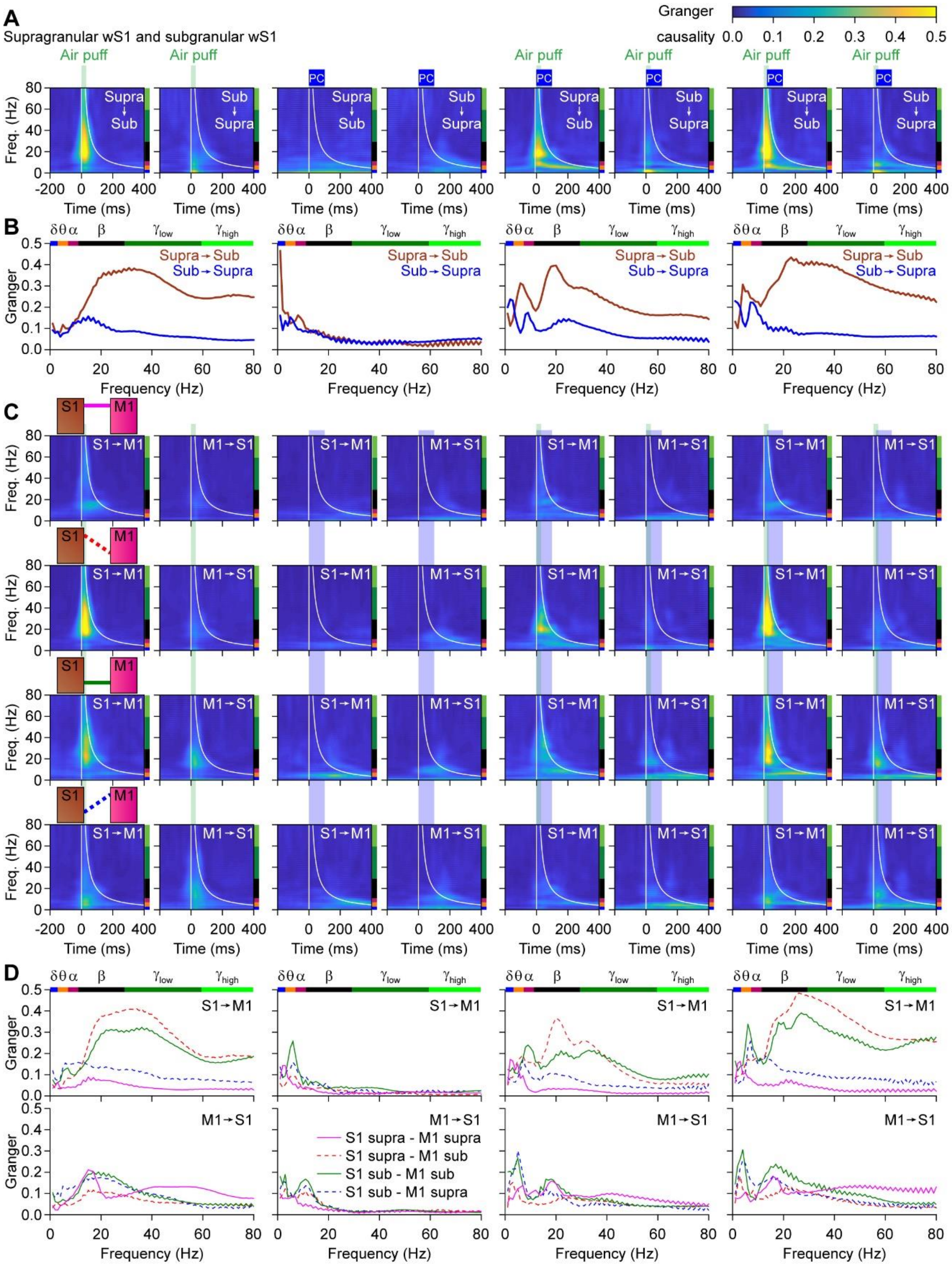


**Fig. S6. Granger causality analysis.**

**A** Granger causality analysis revealed that the sensory-induced beta and gamma band coherence within wS1 was mainly caused by activity in the supragranular layers. The flow from the superficial to the deeper layers in the gamma, but not the beta, band was disrupted by simultaneous Purkinje cell stimulation. **B** The mean Granger causality values for the different conditions, comparing the flow from superficial to deep layers vs. the flow from deep to superficial layers. **C** Granger causality analysis of the coherence between wS1 and wM1, expanding on the analysis shown in Fig. 2C-D where the subgranular layers of wS1 were compared to the supragranular layers of wM1 (data copied in the fourth row to facilitate comparisons). This analysis suggest that the sensory-induced gamma band coherence is mainly caused by the superficial layers of wS1, and from there spreads over wS1 and wM1. The strongest interconnections are found between subgranular layers of wS1 and the supragranular layers of wM1 (fourth row). Also here, wS1 drives wM1 stronger than vice versa, but also wM1 has a share in this coherence, stressing the importance of the connections between wS1 and wM1. **D** The mean Granger causality values for the different relations between wS1 and wM1.

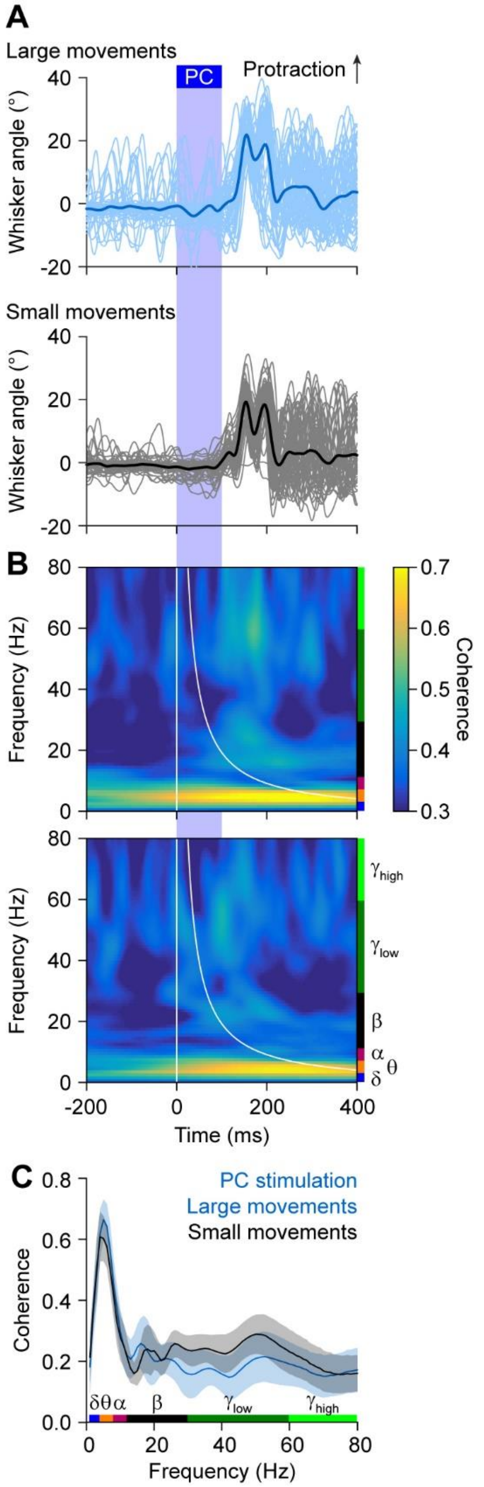


**Fig. S7. Optogenetic Purkinje cell stimulation induces delayed whisker protraction.**
**A** Optogenetic stimulation of Purkinje cells, using a 400 µm diameter optic fiber placed on the center of crus 1, induced whisker protraction at the end of the stimulus. Shown are the 100 trials of a representative experiment, split into the 50% of the trials with the largest and the 50% with the smallest protraction. **B** Heat maps of the coherence over time, showing predominantly activity in the theta band, that was not different between the groups of trials (**C**). *N* = 8 mice. Shades indicate SEM.

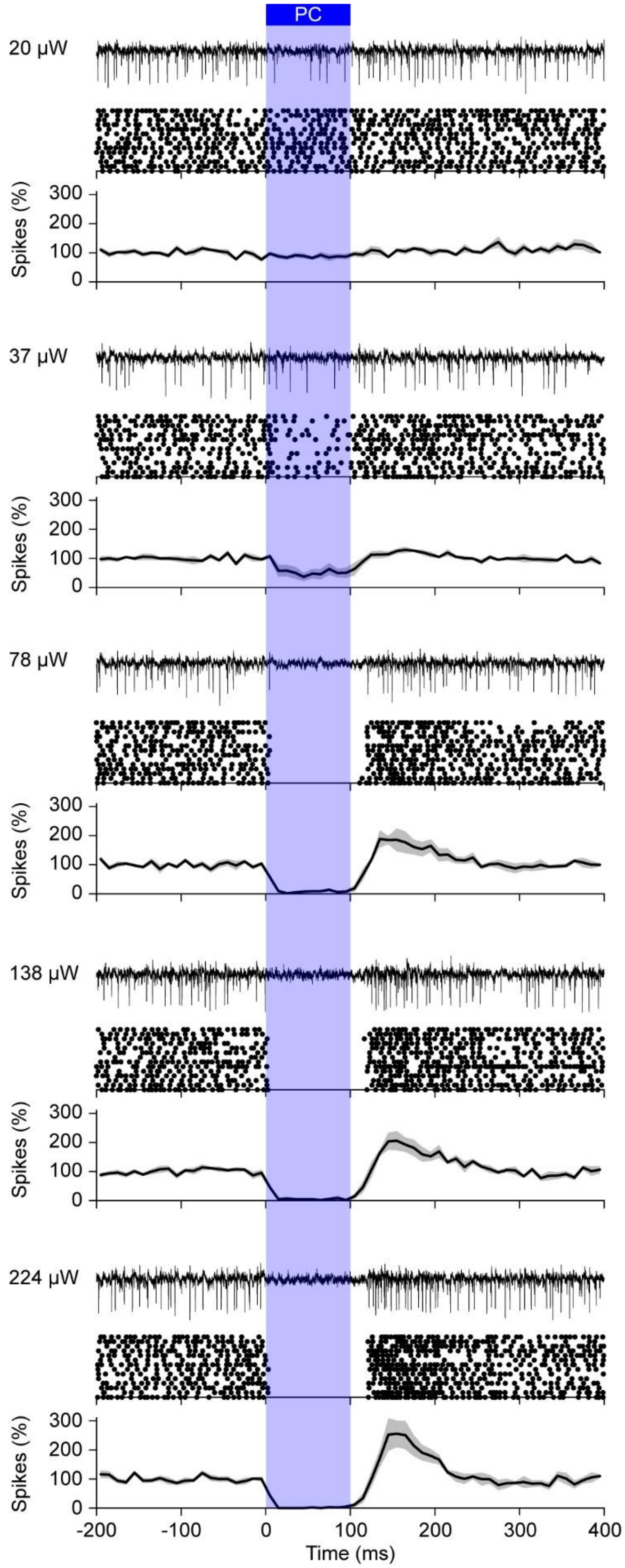


**Fig. S8. Optogenetic stimulation of Purkinje cells silences neurons of the cerebellar nuclei.**

Using different illumination intensities and an optic fiber with a diameter of 105 µm, optogenetic stimulation of Purkinje cells induced a pause in firing of cerebellar nuclei neurons. At higher intensities, the pause was followed by rebound firing. For each intensity, an example trace is plotted, followed by a raster plot of the same experiment and the averaged peri-stimulus histogram of the spike rate (normalized to baseline = 100%) constructed from 6 responsive neurons in *N* = 2 mice. The shades indicate SEM.

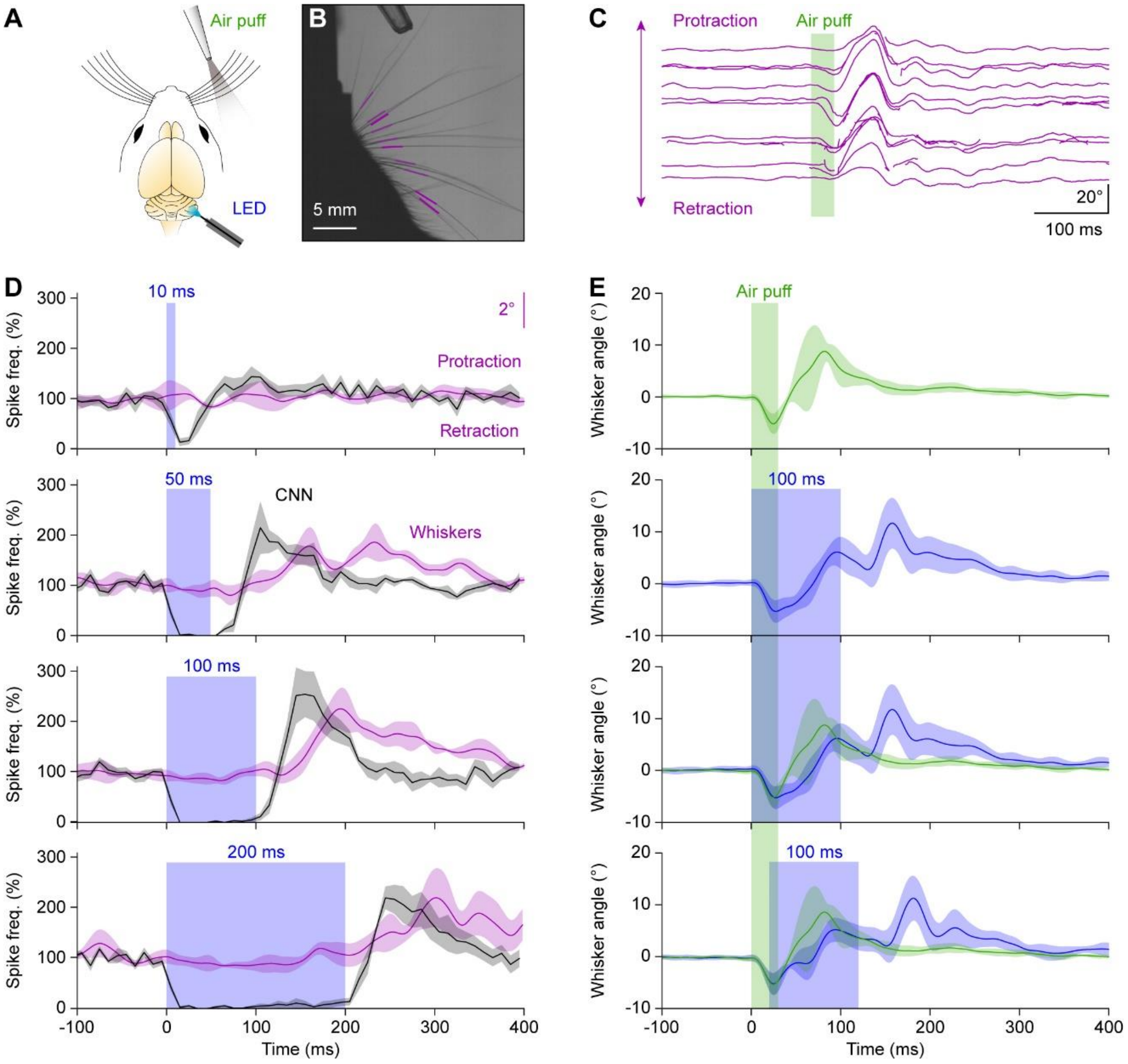


**Fig. S9. Rebound firing in the cerebellar nuclei is linked to whisker protraction.**
**A** Experimental scheme. **B** The movements of the whiskers were tracked using video-analysis. The colored line fragments indicate the tracked part of the whiskers. **C** Raw output of the whisker tracking algorithm, showing for one trial how an air puff blew the whiskers backwards, after which an active protraction followed. **D** Optogenetic stimulation could also trigger whisker protraction, but not during the period of stimulation. By varying the stimulus duration, we observed that the rebound firing in the cerebellar nucleus neurons (CNN) varied in timing and amplitude and that the whisker protraction followed the rebound firing. $n$ = 6 cerebellar nucleus neurons in $N$ = 2 mice. E Whisker air puff stimulation induced a reflexive protraction. This protraction was reduced during the stimulus, but increased at the end of the stimulus. The whisker angle was normalized for each mouse at 0° before stimulus onset. $N$ = 4 mice with each $n$ = 100 trials per condition. Lines indicate average and shades SEM.

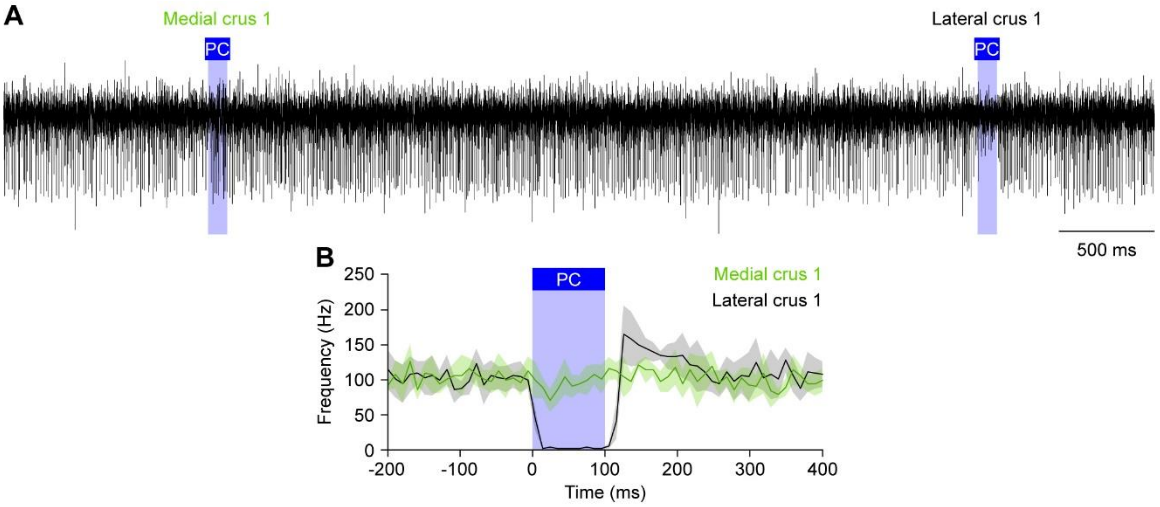


**Fig. S10. Optogenetic Purkinje cell stimulation acts locally.**
**A** Extracellular recording of an exemplary cerebellar nucleus neuron displaying inhibition upon optogenetic stimulation of Purkinje cells in the lateral, but not the medial part of crus 1. For this experiment, an optic fiber with a diameter of 105 µm was used. **B** Averaged peri-stimulus histogram of two simultaneously recorded cerebellar nucleus neurons. The two neurons were separated laterally by 305 µm.

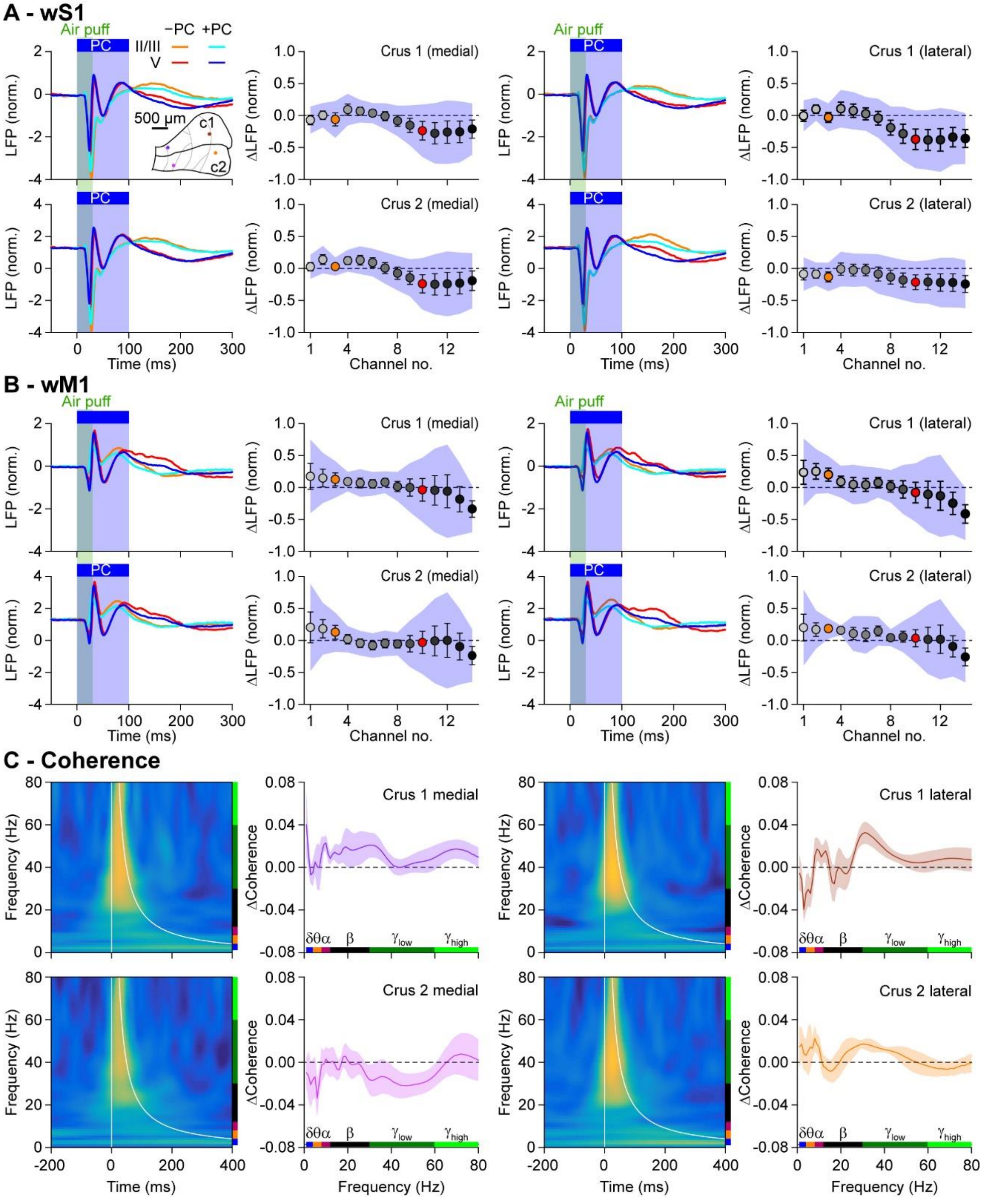


**Fig. S11. Regional differences in the impact of Purkinje cell stimulation on sensory-induced local field potentials in wS1 and wM1.**

**A** Averaged local field potentials (LFP) of the supra- (light colors) and subgranular (dark colors) of wS1 upon either only air puff stimulation of the contralateral facial whiskers (red colors) or air puff stimulation in combination with optogenetic stimulation of Purkinje cells (PC; blue colors). Optogenetic Purkinje cell stimulation was performed using four optic fibers with 105 µm diameter placed at different locations in crus 1 (c1) and crus 2 (c2; inset). During each trial, one of the fibers was activated in a random sequence. The 2nd and 4th columns indicate the difference in the amplitudes of the first positive peaks following stimulation, using the same color codes as in Fig. 1. Error bars indicate SEM and shaded areas sd. **B** The same analysis, but now for wM1. **C** Combined air puff whisker stimulation and optogenetic Purkinje cell stimulation resulted in gamma band coherence between wS1 and wM1 (heat maps). The coherence was different from those trials in which only the whiskers were stimulated, as indicated by the ΔCoherence plots. The impact of Purkinje cell stimulation on sensory-induced coherence depended on the location of optogenetic stimulation, with the lateral part of crus 1 and the medial part of crus 2 having opposite impact on gamma (but not theta) band coherence and the other locations having more intermediate effects. Lines are averages and shaded areas indicate SEM. $N$ = 7 mice.

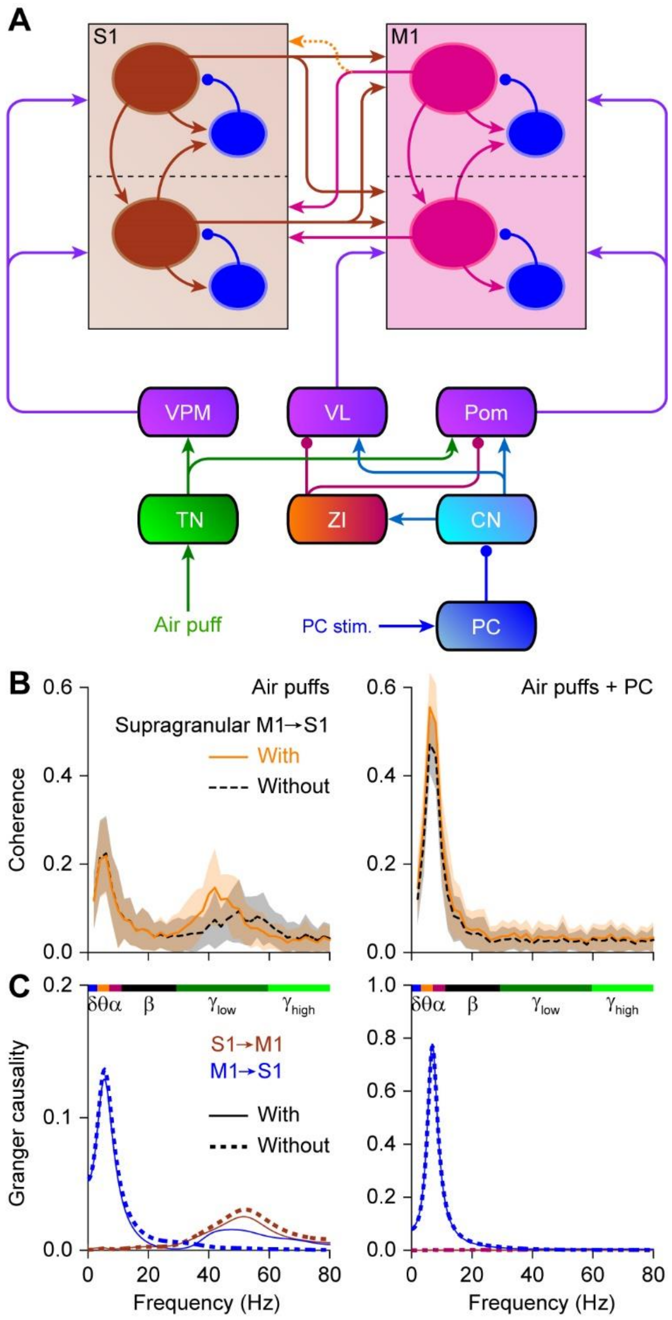


**Fig. S12. Laminar model: impact of a projection for the supragranular layers of M1 to the supragranular layers of S1.**

**A** Here we compared the circuit with and without a direct connection between the supragranular layers of M1 and S1 (dashed orange arrow). **B** Removing the supragranular M1 to S1 connection resulted in a slightly less powerful gamma band coherence (black) upon simulation of the trigeminal nuclei (simulating sensory input of the whiskers) than the same simulation in the presence of the supragranular M1 to S1 connection (orange). The impact of this connection was less during the combined trigeminal + Purkinje cell stimulation. C Granger causality analysis revealed that deleting the supragranular M1 to S1 connection resulted in a virtually complete lack of the contribution of M1 to the sensory-induced gamma band coherence. Note that the situation with the supragranular M1 to S1 connection is the circuit that was used to generate the data of Fig. 5. These data are replicated here to facilitate comparison. Lines indicate averages and shaded areas sd.

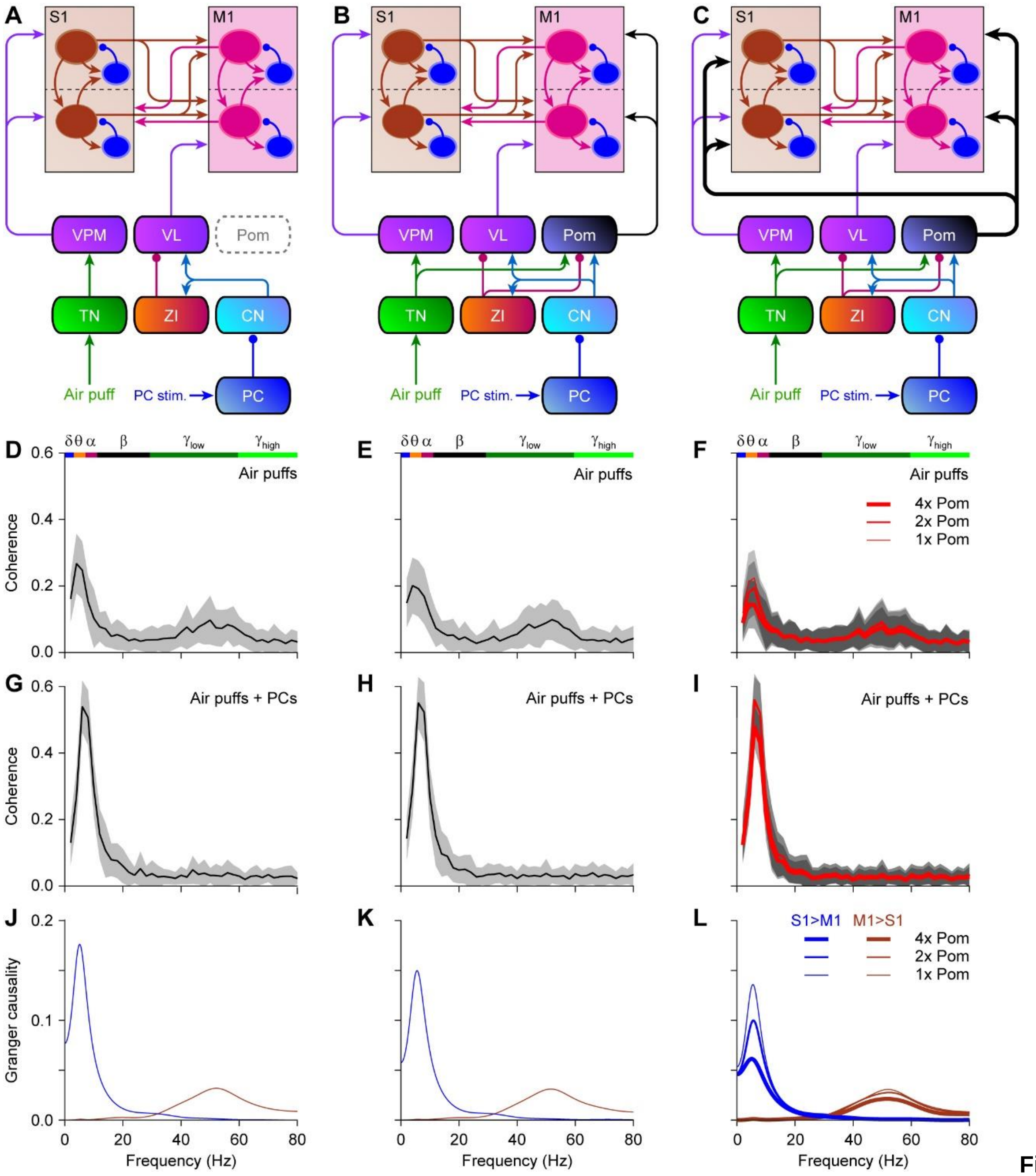


**Fig. S13. Impact of Pom connectivity on laminar model**

To test whether the Pom could affect the flow between S1 and M1 during gamma band coherence, we used our computation model in the configuration without a supragranular connection between M1 and S1 (Fig. S12). In this configuration, S1 is dominant over M1 when generating gamma band coherence. To study the impact of the Pom, we compared three different configurations: without Pom (**A**), with Pom projecting only to M1 (**B**) and with Pom symmetrically projecting to S1 and M1 (**C**). Of the latter, we implemented the connectiivty strength as used for Pom to M1 connection in Fig. 5, double (”2x Pom") and quadruple strength (”4x Pom). **D**-**F** The different configurations did affect the amplitude of sensory-induced coherence between S1 and M1 (simulated by stimulation of the trigeminal nuclei), but did not affect the frequency characteristics. **G**-**I** A similar observation was made for the conjunctive trigeminal + Purkinje cell stimulation. **J**-**L** Granger causality analysis demonstrated that the Pom could not induce M1 to be causative for sensory-induced gamma band coherence (as the direct supragranular connection between M1 and S1 could; see Fig. S12). Lines indicate averages and shaded areas sd.

**Table S1.**

| | ***p*** | $\chi^2$ | Sign.? | Test |
|---|---|---|---|---|
| **First negative peak** | | | | |
| wS1 [supragranular layers] | 0.417 | 1.750 | | Friedman's |
| wS1 [layer IV] | 0.197 | 3.250 | | Friedman's |
| wS1 [subgranular layers] | **0.030** | 7.000 | | Friedman's |
| *Air puff vs. Air puff + simultaneous PC stimulation* | ***0.012*** | *2.500* | *yes* | *Dunn's* |
| *Air puff vs. Air puff + delayed PC stimulation* | *0.046* | *-2.000* | *no* | *Dunn's* |
| *Simultaneous vs. delayed stimulation* | *0.617* | *0.500* | *no* | *Dunn's* |
| wM1 [supragranular layers] | 0.093 | 4.750 | | Friedman's |
| wM1 [subgranular layers] | 0.417 | 1.750 | | Friedman's |
| **First positive peak** | | | | |
| wS1 [supragranular layers] | 0.607 | 1.000 | | Friedman's |
| wS1 [layer IV] | 0.325 | 2.250 | | Friedman's |
| wS1 [subgranular layers] | **0.008** | 9.750 | | Friedman's |
| *Air puff vs. Air puff + simultaneous PC stimulation* | ***0.024*** | *2.250* | *yes* | *Dunn's* |
| *Air puff vs. Air puff + delayed PC stimulation* | *0.453* | *-3.000* | *no* | *Dunn's* |
| *Simultaneous vs. delayed stimulation* | ***0.003*** | *-0.750* | *yes* | *Dunn's* |
| wM1 [subgranular layers] | **0.030** | 7.000 | | Friedman's |
| *Air puff vs. Air puff + simultaneous PC stimulation* | 0.617 | -0.500 | *no* | *Dunn's* |
| *Air puff vs. Air puff + delayed PC stimulation* | **0.012** | -2.500 | *yes* | *Dunn's* |
| *Simultaneous vs. delayed stimulation* | 0.046 | -2.000 | *no* | *Dunn's* |
| wM1 [subgranular layers] | 0.008 | 9.750 | | Friedman's |
| *Air puff vs. Air puff + simultaneous PC stimulation* | ***0.024*** | *2.250* | *no* | *Dunn's* |
| *Air puff vs. Air puff + delayed PC stimulation* | *0.453* | *-3.000* | *yes* | *Dunn's* |
| *Simultaneous vs. delayed stimulation* | ***0.003*** | *-0.750* | yes | *Dunn's* |

**Statistical evaluation of the data represented in Figs. 1 and S4.** For each mouse, the averages of the first negative and the first positive peak were compared between three conditions (only air puff stimulation of the whiskers, simultaneous air puff and optogenetic Purkinje cell (PC) stimulation and air puff stimulation combined with a 20 ms delayed PC stimulation. Averages were compared with Friedman's two-way ANOVA and, if significant, with pair-wise Dunn's post-tests. The *p* values of the post-tests were not corrected for multiple comparisons, but Benjamini-Hochberg correction was performed for multiple comparisons among post-tests and the outcomes are listed as statistically significant or not in the column "Sign.?".

#### SI References


1. J. Potworowski, W. Jakuczun, S. Leski, D. Wójcik, Kernel current source density method. *Neural Computation* **24**, 541-575 (2012).
2. R. Oostenveld, P. Fries, E. Maris, J. M. Schoffelen, FieldTrip: Open source software for advanced analysis of MEG, EEG, and invasive electrophysiological data. *Comput Intell Neurosci* **2011**, 156869 (2011).
3. A. M. Amjad, D. M. Halliday, J. R. Rosenberg, B. A. Conway, An extended difference of coherence test for comparing and combining several independent coherence estimates: theory and application to the study of motor units and physiological tremor. *J Neurosci Methods* **73**, 69-79 (1997).
4. I. Perkon, A. Kosir, P. M. Itskov, J. Tasic, M. E. Diamond, Unsupervised quantification of whisking and head movement in freely moving rodents. *J Neurophysiol* **105**, 1950-1962 (2011).
5. N. Rahmati *et al.*, Cerebellar potentiation and learning a whisker-based object localization task with a time response window. *J Neurosci* **34**, 1949-1962 (2014).
6. V. Romano *et al.*, Potentiation of cerebellar Purkinje cells facilitates whisker reflex adaptation through increased simple spike activity. *eLife* **7**, e38852 (2018).
